# Breakdown of spatial coding and neural synchronization in epilepsy

**DOI:** 10.1101/358580

**Authors:** Tristan Shuman, Daniel Aharoni, Denise J. Cai, Christopher R. Lee, Spyridon Chavlis, Jiannis Taxidis, Sergio E. Flores, Kevin Cheng, Milad Javaherian, Christina C. Kaba, Matthew Shtrahman, Konstantin I. Bakhurin, Sotiris Masmanidis, Baljit S. Khakh, Panayiota Poirazi, Alcino J. Silva, Peyman Golshani

## Abstract

Temporal lobe epilepsy causes significant cognitive deficits in both human patients and rodent models, yet the specific circuit mechanisms that alter cognitive processes remain unknown. There is dramatic and selective interneuron death and axonal reorganization within the hippocampus of both humans and animal models, but the functional consequences of these changes on information processing at the neuronal population level have not been well characterized. To examine spatial representations of epileptic and control mice, we developed a novel wire-free miniature microscope to allow for unconstrained behavior during *in vivo* calcium imaging of neuronal activity. We found that epileptic mice running on a linear track had severely impaired spatial processing in CA1 within a single session, as place cells were less precise and less stable, and population coding was impaired. Long-term stability of place cells was also compromised as place cells in epileptic mice were highly unstable across short time intervals and completely remapped across a week. Because of the large-scale reorganization of inhibitory circuits in epilepsy, we hypothesized that degraded spatial representations were caused by dysfunctional inhibition. To test this hypothesis, we examined the temporal dynamics of hippocampal interneurons using silicon probes to simultaneously record from CA1 and dentate gyrus during head-fixed virtual navigation. We found that epileptic mice had a profound reduction in theta coherence between the dentate gyrus and CA1 regions and altered interneuron synchronization. In particular, dentate interneurons of epileptic mice had altered phase preferences to ongoing theta oscillations, which decorrelated inhibitory population firing between CA1 and dentate gyrus. To assess the specific contribution of desynchronization on spatial coding, we built a CA1 network model to simulate hippocampal desynchronization. Critically, we found that desynchronized inputs reduced the information content and stability of CA1 neurons, consistent with the experimental data. Together, these results demonstrate that temporally precise intra-hippocampal communication is critical for forming the spatial code and that desynchronized firing of hippocampal neuronal populations contributes to poor spatial processing in epileptic mice.

## Main Text

Temporal lobe epilepsy (TLE) is associated with disabling cognitive deficits^1,2^, interneuron cell death^3,4^, and large-scale anatomical reorganization of limbic circuits^5–8^ in both human patients and rodent models. Following cell death, surviving interneurons from both CA1 and dentate gyrus (DG) sprout new local and long-range connections leading to altered timing and kinetics of inhibition^8–12^. These alterations are likely to have dramatic effects on spatial processing as the hippocampus relies on the precise timing of diverse interneuron subtypes that control excitatory inputs^13–16^. Indeed, initial studies have shown degraded spatial representations in epileptic rodents^17–20^, but it remains unclear whether chronic epilepsy disrupts the formation or maintenance of spatial maps, how population coding is affected, and how stable these representations are across long time spans. To address these questions, we used *in vivo* calcium imaging with miniature microscopes to track large numbers of neurons during spatial navigation across many days and to characterize the specific properties of spatial processing that are altered in epileptic mice.

Recent developments in miniature microscope technology have allowed unprecedented access to neural signals in freely behaving animals^21–24^, yet all commercially available miniature microscopes use tethered cables to transmit power and data to a recording acquisition device. In order to allow for more naturalistic behaviors and improve behavioral performance during recordings, we developed an open-source wire-free miniature microscope that allows us to perform calcium imaging without a cable tethered to the animal’s head (Fig 1A, Extended Data 1A,B). This wire-free Miniscope saves data directly to a microSD card positioned on the Miniscope and is powered by a lithium-polymer battery mounted to its side. Together, this configuration allows for completely untethered and more naturalistic behavior during neural recording (Extended Data 1C,D).

**Figure 1.**
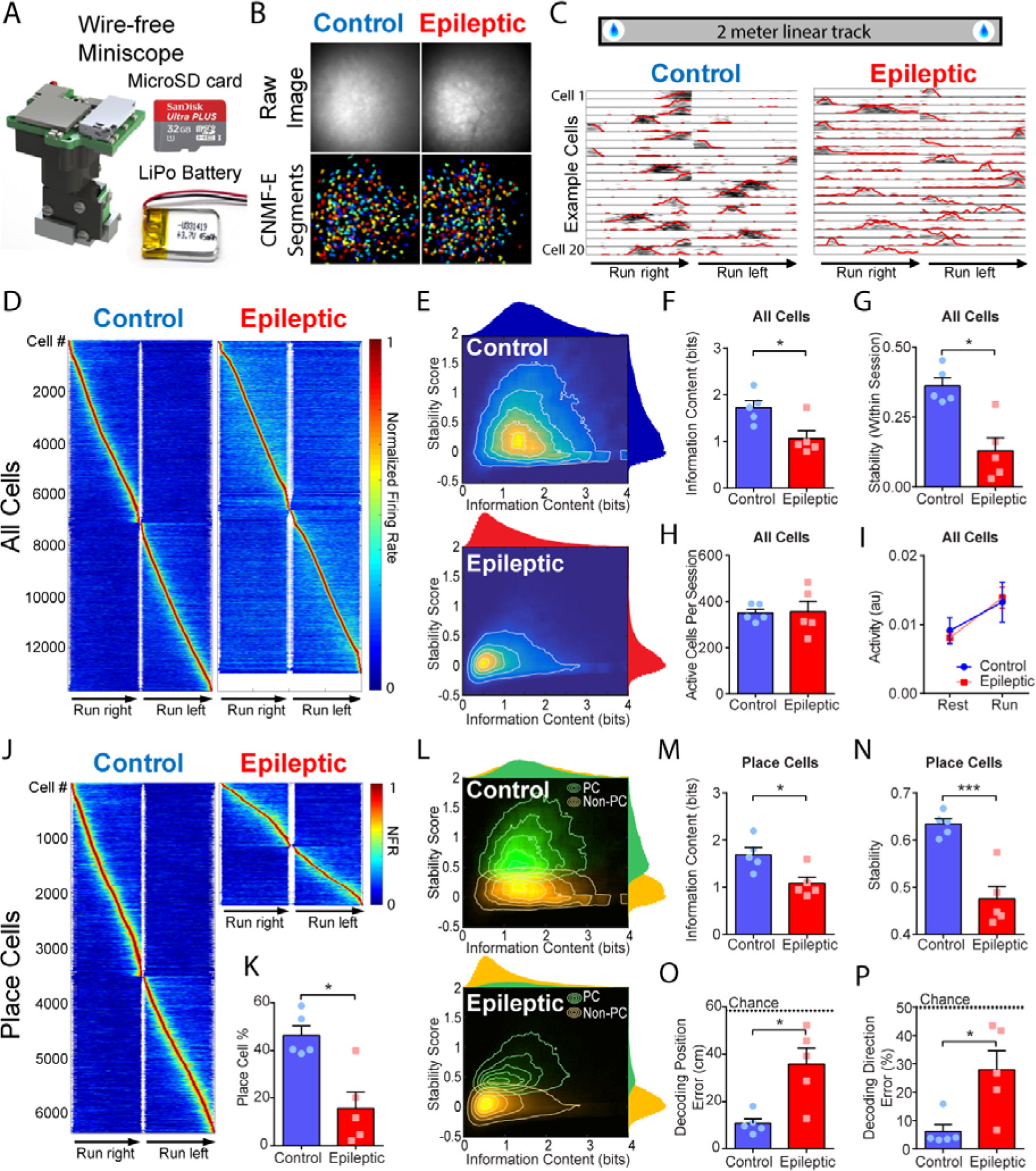
Disrupted spatial coding in epileptic mice. A. Wire-free Miniscope for untethered behavior during imaging. The miniscope is powered by a lithium-polymer battery and data is saved directly to a microSD card. B. Example raw images (top) and identified cells (bottom) in control and epileptic mice. Each field of view is approximately 550um x 550um. Cells were identified with CNMF-E. C. Animals were trained to run on a 2m linear track for water reward. Bottom, twenty example cells from control and epileptic mice show spatially selective firing patterns. D. Normalized spatial firing rates across all cells recorded during all sessions (4-8 sessions per animal). E. Heatmap of stability and information content in control (top) and epileptic (bottom) mice. Contours represent 10^th^, 30^th^, 50^th^, 70^th^, and 90^th^ percentiles. Note the shift in the distributions of epileptic mice toward lower information content and stability. F. Average information content across all cells for each animal. Epileptic mice had less information content than control mice (Mann-Whitney test, P=0.03). G. Average stability across all cells for each animal. Epileptic mice had less stable cells than control mice (Unpaired t-test, P=0.02). H. Average number of active cells per session. No differences in the number of active cells were found (Unpaired t-test, P=0.73). I. Average activity of all cells during run and rest. No differences in activity were found (2-way ANOVA, F_Group_ (1,8)=0.01, P=0.94, Run vs Rest: P<0.05 for both groups). J. Normalized spatial firing rates across place cells (see Methods) recorded during all sessions (4-8 sessions per animal). K. Average percent of all cells that were place cells during each session. Epileptic mice had less place cells than control mice (Unpaired t-test, P=0.03). L. Heatmap of stability and information content in control (top) and epileptic (bottom) mice in both place cells (green) and non-place cells (yellow). Contours represent 10^th^, 30 ^th^, 50 ^th^, 70 ^th^, and 90 ^th^ percentiles. M. Average information content across place cells for each animal. Place cells in epileptic mice had less information content than in control mice (Unpaired t-test, P=0.02). N. Average stability across place cells for each animal. Place cells in epileptic mice were less stable than in control mice (Unpaired t-test, P=0.001). O. Average decoding error for position within each session. Epileptic mice had larger decoding position error than control mice (Unpaired t-test, P=0.03). P. Average decoding error for direction within each session. Epileptic mice had more decoding position error than control mice (Mann-Whitney test, P=0.03). N=5 per group for all panels. Error bars represent 1 S.E.M. *P<0.05.

To study spatial processing in a mouse model of TLE, we used an established model where an initial prolonged seizure induces chronic epilepsy and severe spatial memory deficits for the life of the animal^2^. Naïve mice were injected with pilocarpine to induce a 2-hr status epilepticus event. All animals that recovered displayed spontaneous seizures and cognitive deficits at least 6 weeks later (Extended Data 2). To record place cell activity, AAV1-Syn-GCaMP6f virus was injected into CA1 of dorsal hippocampus at least 4 weeks after status epilepticus. A GRIN lens was then implanted above CA1 and mice were trained to run on a 2 meter linear track for water reward while wearing a wire-free Miniscope (Fig 1B, C, Supplementary Video 1). Within a single session, both epileptic and control mice had spatially tuned firing patterns along the track (Fig 1D). However, the quality of the spatial information was drastically reduced in the epileptic mice. We measured the information content (how specific each neuron codes for spatial location) and stability (how reliably each neuron fires at that spatial location across trials) of all cells and found a dramatic reduction of both measures in epileptic mice (Fig 1E-G). In addition, the population vector overlap between locations along the track was significantly higher in the epileptic group (Extended Data 3A,B) indicating that neuronal populations are less tuned for spatial position. These effects were not due to differences in the overall number of cells active during each session or the activity level of neurons during running or rest (Fig 1 H,I). We next separated the neuronal population into place cells and non-place cells according to their information content and stability (see Methods) and tested whether the quality of place cells, specifically, was diminished in epileptic mice (Fig 1J). Epileptic mice had fewer place cells as a percent of total active cells (Fig 1K). In addition, the place cells of epileptic mice had significantly lower information content and stability within a session (Fig 1L-N), as well as increased population vector overlap between different locations the track (Extended Data 3C,D). While a linear decoder could accurately predict the position of the control mouse on interleaved trials, we found that the decoding accuracy was significantly impaired in the epileptic mice for both position (Fig 1O) and running direction (Fig 1P). Together, these findings illustrate a severely disrupted spatial code in the epileptic hippocampus and suggest that cognitive deficits in epilepsy may be due to the inability to create and maintain consistent spatial information.

Next we examined the long-term stability of place cells across many days. We imaged animals running on the same linear track with varying inter-session intervals at 30 minutes, 6 hours, 24 hours, 2 days and 7 days (Fig 2A-C, Extended Data 4). In epileptic mice, the spatial firing of a large proportion of the place cells dramatically remapped when epileptic mice returned to the same track 30 minutes later, while those in control mice remained stable (Fig 2A). Across 7 days, in control mice, a large proportion of place cells continued to fire at the same location, while in epileptic animals very few neurons fired at similar locations. (Fig 2B). This instability across sessions is evident in the stability of individual place cells (Fig 2D), as well as at the population level as epileptic mice had higher PVO between large (>66cm) offset distances across trials (Fig 2C). In addition, place cells in epileptic mice were less likely to remain place cells in subsequent sessions (Fig 2E). This was not due to turnover in the active population as cells were equally likely to be active in subsequent sessions in the epileptic mice (Fig 2F), even when only considering place cells (Fig 2G). The instability of place cells led to significantly worse decoding of position and running direction across all time intervals in epileptic animals (Fig 2H,I, Supplementary Video 2). Together, these data demonstrate that epileptic mice have unstable spatial representations, even across 30 minutes, which continue to degrade to chance levels within a week. Furthermore, these results provide additional evidence that deficits in initially establishing stable spatial information are a primary contributor to place cell instability in epileptic mice.

**Figure 2.**
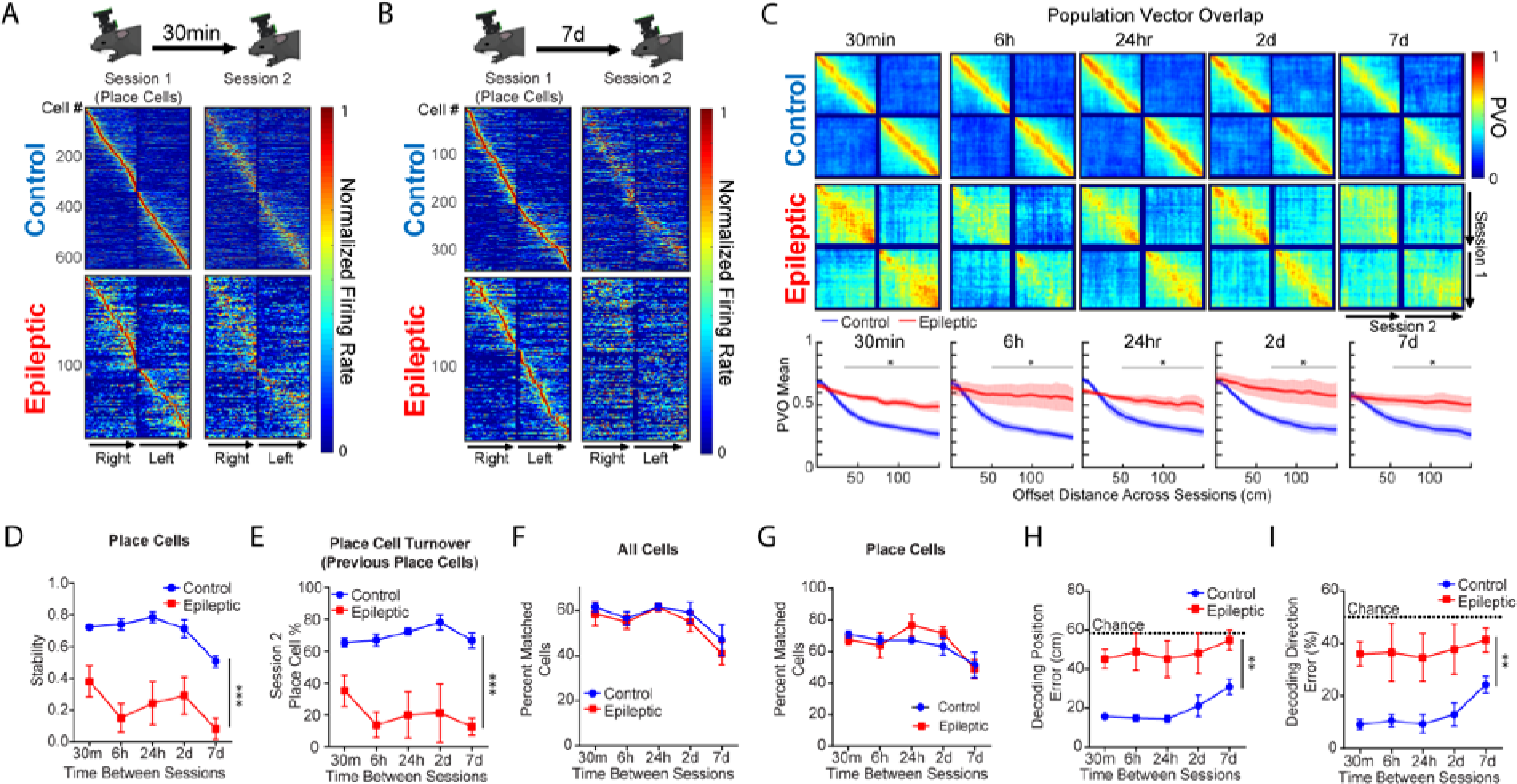
Disrupted stability of place cells across sessions. A,B. Animals ran on the same linear track with 30min, 6h, 24h, 2d, or 7d between sessions. Normalized spatial firing rates of place cells on session 1 that were also active on the second session 30 minutes (A) or 7 days (B) later. Cells are sorted by the peak firing rate on the first session only. C. Top, Population vector overlap (PVO) across sessions for place cells recorded at each time interval. Bottom, Mean PVO across animals. For all time points, Epileptic mice had higher PVO (less distinct firing patterns) at offset distances above 34-66cm (30min: P<0.05 for 34-150cm, 6h: P<0.05 for 50-150cm, 24hr: P<0.05 for 50-150cm, 2d: P<0.05 for 66-150cm, 7d: P<0.05 for 52-150cm). D. Stability across sessions at each time interval. Epileptic mice had lower stability across all time points (2-way ANOVA, F_Group_ (1,7.75)=26.75, P=0.001, post hoc for all time points: P<0.05). E. Place cell percentage of cells that were place cells on session 1. Epileptic mice had less place cells across all time intervals (2-way ANOVA, F_Group_(1,8.27)=34.8, P<0.001, post hoc for all time points: P<0.05). F. Percent of all cells active during session 1 that were active again during session 2 at each time interval. There were no differences between groups (2-way ANOVA, F_Group_(1,4.2)=0.008, P=0.93). G. Percent of place cells active during session 1 that were active again during session 2 at each time interval. There were no differences between groups (2-way ANOVA, F_Group_(1,5.48)=0.635, P=0.46). H. Decoding position error across sessions. Epileptic mice had higher error across all intervals (2-way ANOVA, F_Group_(1,8.11)=17.85, P=0.003, post hoc for all time points: P<0.05). I. Decoding direction error across sessions. Epileptic mice had higher error across all intervals (2-way ANOVA, F_Group_(1,8.13)=11.57, P=0.009, post hoc for all time points: P<0.05). N=5 per group for all panels. Error bars represent 1 S.E.M. *P<0.05, **P<0.01, ***P<0.001.

A predominant feature of the epileptic brain in both patients and mouse models is substantial interneuron death and axonal reorganization^3–8^. In particular, in chronically epileptic mice, new functional inhibitory connections form within the DG^6^, between CA1 and DG^7^, and even across hemispheres^25^. Concurrently, network activity is altered in the hippocampus of epileptic mice, with reductions in the power of theta oscillations that correlate with memory impairments^26–28^. In addition, changes in the synchrony of hippocampal rhythms have been hypothesized to underlie cognitive deficits^28,29^. Since interneurons are known to control information processing in the hippocampus^13–16^, we hypothesized that reorganization of the hippocampal network would lead to altered synchronization of inhibition in the hippocampus of epileptic mice. To test this hypothesis, we performed silicon probe recordings in head-fixed epileptic and control mice running on a linear track in virtual reality (Fig 3A,B, Supplementary Video 3). Following confirmation of chronic epilepsy, we implanted a head-bar and trained the mice to run through a virtual linear track in order to record hippocampal activity during active spatial processing (see Methods). Both groups learned to run through the virtual linear track and there were no differences in the number of trials completed, time spent running, or running speed (Extended Data 5D-F). After animals were trained to complete hundreds of trials within a single session, we performed acute extracellular electrophysiology with 128 channel silicon probes that spanned the layers of CA1 and DG (Fig 3B,C).

**Figure 3.**
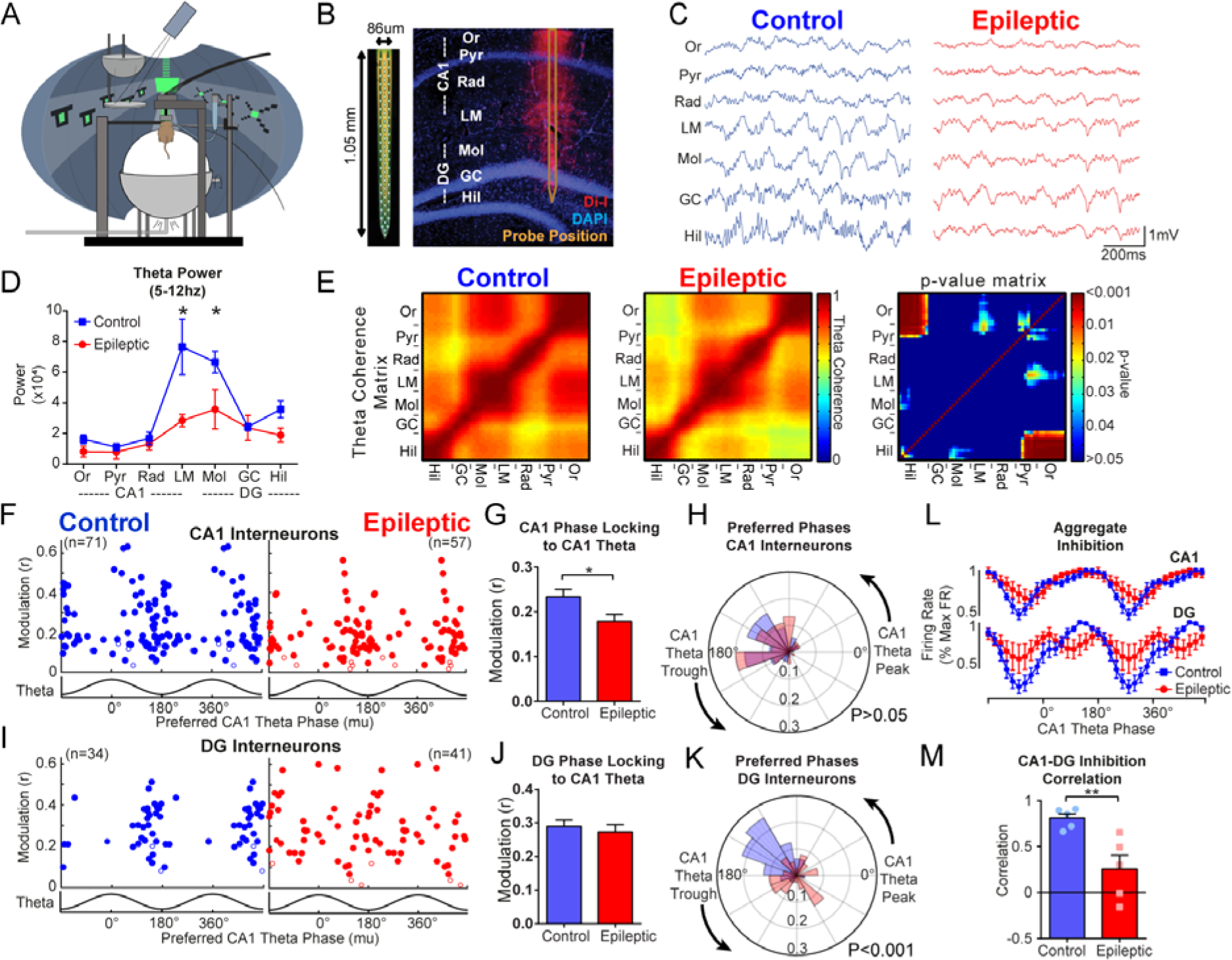
Desynchronization of hippocampal inhibition in epileptic mice. A. Head-fixed mice were trained to run through a virtual linear track for water rewards. B. During virtual navigation, silicon probes were inserted into the dorsal hippocampus spanning the CA1 and DG. C. Example data from each layer of the hippocampus during locomotion. D. Theta power from each layer. Epileptic mice had reduced power in the lacunosum moleculare and molecular layer of DG (FGroupXRegion(6,48)=5.30, P<0.001, LM: P<0.001, Mol: P<0.05). E. Theta coherence between each channel pair along the recording probe in control (left) and epileptic (middle) mice. Right, p-value matrix for each pair of recording sites (Unpaired t-test for each channel pair). F. Phase locking to CA1 theta for each interneuron in CA1 of control and epileptic mice (See Extended Data 7). Each dot represents one interneuron and the data is double plotted for visualization. Closed circles were significantly phase locked (Raleigh test) and open circles were not. G. Mean phase modulation (r) of CA1 interneurons to CA1 theta. Interneurons in epileptic mice had reduced phase modulation (n=71 Control cells, n=57 Epileptic cells, Mann-Whitney test, P=0.02). H. Rose plot of preferred firing phases of significantly phase locked CA1 interneurons. There were no differences between interneurons in control and epileptic mice (Kuiper circular test, P>0.05). I. Phase locking to CA1 theta for each interneuron in DG of control and epileptic mice. J. Mean phase modulation (r) of DG interneurons to CA1 theta. No difference was found between control and epileptic interneurons (n=34 Control cells, n=41 Epileptic cells, Unpaired t-test, P=0.57). K. Rose plot of preferred firing phases of significantly phase locked DG interneurons. The distribution of preferred phases in control and epileptic mice were different (Kuiper circular test, P<0.001). L. Aggregate inhibition of all interneurons in CA1 and DG relative to CA1 theta phase. No differences were found in CA1 (F_GroupXPhase_(17,136)=1.526, P=0.09) but a significant interaction was found in DG (F_GroupXPhase_(17,136)=3.089, P<0.001) indicating abnormal aggregate inhibition in DG. M. Correlation between CA1 and DG aggregate inhibition. Epileptic mice had lower correlation between CA1 and DG aggregate firing rates than control mice (Unpaired t-test, P<0.01). Or, stratum oriens; Pyr, stratum pyramidale; Rad, stratum radiatum; LM, stratum lacunosum moleculare; Mol, molecular layer of dentate gyrus; GC, granule cell layer; Hil, hilus. N=5 animals per group for all panels. Error bars represent 1 S.E.M. *P<0.05, **P<0.01

We first examined the power of local field potentials during locomotion throughout CA1 and DG across several frequencies (Extended Data 6, 8). In epileptic mice, similar to previous reports^26–28^, there was significantly reduced theta power in the lacunosum moleculare layer of CA1 and molecular layer of DG (Fig 3D). To examine the synchrony of theta oscillations along the CA1-DG axis, we measured phase coherence of theta oscillations between each electrode pair and found reduced theta coherence primarily between the pyramidal layer and stratum oriens of CA1 and the dentate hilus (Fig 3E). This reduced theta power and coherence in the epileptic hippocampus suggests a decrease in the synchronization of excitatory and inhibitory synaptic inputs onto the two structures, which may arise from desynchronized firing of interneurons in CA1 and DG. To test this hypothesis, we examined the coordinated firing patterns of interneurons in both CA1 and DG. As previously reported^13,14^, individual CA1 interneurons modulate their firing rates in phase with theta oscillations (Extended Data 7), which can be measured with a modulation index (r) and preferred phase (mu). Similar to previous studies, we found that CA1 interneurons in control animals were highly phase locked to the falling phase and trough of CA1 theta (Fig 3F, left). In epileptic mice, CA1 interneurons had reduced phase locking (Fig 3F,G) but maintained a similar preferred phase to control mice (Fig 3F,H). This reduction in phase locking was not solely due to reduced theta power as the effect was maintained when the power was matched across groups (Extended Data 9A-D). In the DG of control mice, interneurons were almost exclusively phase locked near the falling phase and trough of CA1 theta (Fig 3I), indicating high synchronization of interneurons across the hippocampal subregions. In the epileptic mice, individual interneurons had equivalent magnitude of phase locking (Fig 3J), but the preferred phases of all the neurons were highly distributed across all phases of CA1 theta (Fig 3I,K). This observation was maintained when referenced to theta in lacunosum moleculare or the dentate hilus (Extended Data 9E-H). Thus, in control animals the aggregate inhibition in CA1 and DG is highly synchronous and peaks near the trough of CA1 theta (Fig 3L). In the epileptic mice, the DG inhibition is distributed across all phases of theta (Fig 3L), which desynchronizes the two populations and leads to a reduced correlation between total inhibition in CA1 and DG (Fig 3M). Together, these results indicate interneuron firing in CA1 and DG regions become desynchronized in epileptic mice, which likely contributes to the profound spatial processing deficits we have found.

Our electrophysiological findings demonstrate that synchronization mechanisms in the epileptic hippocampus are severely impaired, with abnormal firing patterns in this circuit critical for spatial coding. To test how hippocampal desynchronization specifically impairs spatial processing, we built a CA1 network model to simulate place cell processing from grid-like inputs (Fig 4A,B). This model provides temporally offset inputs from entorhinal cortex and DG-CA3 pathways with feedforward and feedback inhibition from several interneuron subtypes (Fig 4B) and produces place cell outputs from CA1 pyramidal neurons (Fig 4C). We tested the effects of removing different subclasses of interneurons (mimicking changes observed in epilepsy^9,10^) as well as desynchronizing the inputs into CA1, and measured the spatial output of pyramidal neurons (Fig 4C). We found that removing parvalbumin-positive (PV+), somatostatin-positive (SOM+), or both PV+ and SOM+ interneurons had no effect on the number of place cells, spatial information, or stability of CA1 place cells (Fig 4D-H). In contrast, desynchronization of inputs significantly reduced the number of place cells as well as the spatial information and stability of these place cells (Fig 4D-H), recapitulating our experimental findings. Together, these results indicate that desynchronization of inputs is the primary driver of altered spatial information processing in our model and likely contributes to poor spatial processing in temporal lobe epilepsy.

**Figure 4.**
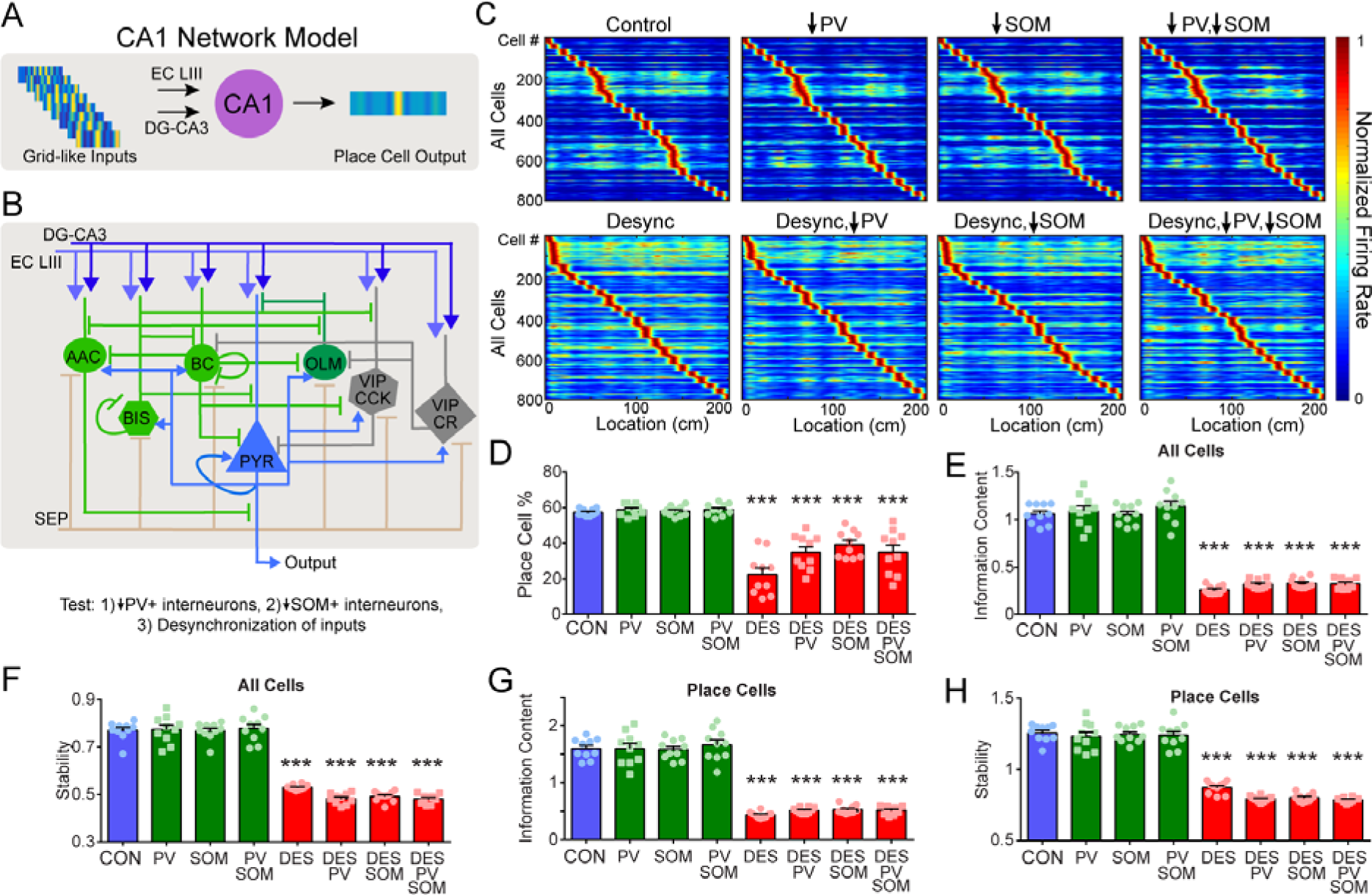
Desynchronization disrupts place cell coding in a CA1 network model. A. Schematic of CA1 network model. Grid-like inputs from EC and DG-CA3 are processed by CA1 and output place cell representations. B. Detailed model including temporally offset excitatory inputs, PV+ interneurons (AXO, BC, BIS), SOM+ interneurons (OLM), and VIP interneurons (VIP/CCK, VIP/CR). We tested the effects of decreasing PV+ interneurons, SOM+ interneurons, and desynchronization, which are all present in chronic epilepsy. C. Normalized spatial firing rates for each conditioned tested. D. Percent of excitatory cells that were place cells. All groups with desynchronization had reduced place cells, while decreasing PV+, SOM+, or both types of interneurons did not (F(7,72)=35.06, P<0.001). E. For all cells, information content was reduced only in the conditions with desynchronization (F(7,72)=156.5, P<0.001) F. For all cells, stability was reduced only in the conditions with desynchronization (F(7,72)=76.3, P<0.001). G. For place cells, information content was reduced only in the conditions with desynchronization (F(7,72)=122.3, P<0.001). H. For place cells, stability was reduced only in the conditions with desynchronization (F(7,72)=139.6, P<0.001). N=10 simulated animals with 80 pyramidal cells per group for all panels. AAC, axo-axonic cells; BC, basket cells; BIS, bistratified cells; OLM, oriens-lacunosum moleculare cells; VIP-CCK, vasoactive intestinal peptide-cholecystokinin cells; VIP-CR, vasoactive intestinal peptide-calretinin. Error bars represent 1 S.E.M. ***P<0.001, all comparisons against the Control condition.

Memory deficits are a core feature of temporal lobe epilepsy and understanding how cognitive impairments arise is a major priority for the epilepsy field^30^. Our results are consistent with prior findings showing poor spatial processing in models of epilepsy^17–20,26–29^ and suggest a novel mechanism causing these spatial deficits independent from seizure activity. Spatial memory relies on precise timing of interneurons that shape excitatory information processing through the hippocampus. We have now discovered that in epileptic mice the timing of inhibitory firing is disrupted and without properly timed excitation and inhibition, this circuit is unable to appropriately process spatial information. In future studies, it will be critical to identify the specific mechanisms that desynchronize this circuit in order to precisely target these mechanisms with new therapeutics for treating cognitive deficits associated with epilepsy.

## Methods

**Subjects:** All experimental protocols were approved by the Chancellor’s Animal Research Committee of the University of California, Los Angeles, in accordance with NIH guidelines. Seven week old male C57B6 mice from Charles River Laboratories were group housed (2-5 per cage) on a 12 h light/dark cycle for all experiments.

**Viral construct:** AAV1.Syn.GCaMP6f.WPRE.SV40 virus (titre: 4.65 × 10^13^ GC per ml) was purchased from Penn Vector Core.

**Epileptogenesis and monitoring:** 7 week old mice were randomly assigned to receive pilocarpine or control injections. All mice first received 1mg/kg i.p. injections of scopolamine methyl bromide (Sigma, CAS 155-41-9) to reduce peripheral effects of pilocarpine. Thirty minutes later, pilocarpine-treated mice received 275mg/kg i.p. injections of pilocarpine hydrochloride (Sigma, CAS 54-71-7) and all animals were monitored continuously for seizures. After 45 minutes, pilocarpine-treated mice that did not show behavioral seizures received additional pilocarpine injections of 50-100mg/kg. The beginning of status epilepticus was recorded as the time of the final behavioral seizure prior to entering continuous seizure activity. After 2hr of status epilepticus, all mice received 10mg/kg i.p. injections of diazepam and were closely monitored for 48 hours to ensure recovery. Pilocarpine-treated mice that did not enter status epilepticus or showed continued negative health effects 48 hours after pilocarpine were removed from the experiment. To confirm chronic epilepsy, mice were video monitored in their home cage and seizures were recorded by a blind observer. In all experiments, spontaneous seizures were confirmed in all epileptic mice.

**Morris water maze:** Morris water maze^31^ was performed with a circular tub filled with a combination of water and white paint. For training, a hidden platform was submerged in one quadrant and start locations were randomized to one of four equidistant locations. Animals were trained three times per day for six days. In each trial, mice were given 60 s to find the platform. If mice found the platform earlier than 60 s, the trial ended then. If mice failed to find the platform, the trial terminated at 60 s. After each trial, mice were put on the platform for 15 s. One day after the last training session, animals were given a probe trial with the platform removed and animals were allowed to swim freely for 60 seconds. Animal location was tracked by an overhead camera running Actimetrics software.

**Cued, delayed alternation T-maze:** T-maze experiments were performed similar to previously described^32^. Prior to the experiment, mice were water restricted and maintained at a body weight of ~85% of initial weight. Mice were handled for three minutes per day for five days and were then trained for 5 days. On each training day, mice received up to 20 trials within a limit of 45 minutes. Each trial consisted of 3 phases: Cue Phase, Delay Phase, and Test Phase (see Extended Data 2B). During the Cue Phase, animals began at the end of the long arm and one side of the T-maze was closed and one side remained open. Animals ran to the open arm to drink the 10uL water reward, and then ran back to the starting point and received a second 10uL water reward. During the Delay Phase, animals were then confined to the starting location for 15 seconds while the maze was cleaned with 7% ethanol. For the Test Phase, the animal was released and allowed to choose one arm to visit. If the animal chose the side opposite the cue, they received a 10uL water reward. If they chose the same side as the cue, they did not receive a reward. They then ran back to the starting point, received another water reward and were confined to the start zone until the next trial began 30 seconds later.

**Calcium imaging surgeries:** For all surgeries, mice were anaesthetized with 1.5 to 2.0% isoflurane and placed into a stereotactic frame (David Kopf Instruments, Tujunga, CA). Lidocaine (2%; Akorn, Lake Forest, Illinois) was applied to the sterilized incision site as an analgesic, while subcutaneous saline injections were administered throughout each surgical procedure to prevent dehydration.

For calcium imaging experiments, all mice underwent two stereotaxic surgeries. First, mice were unilaterally injected with 500nl of AAV1.Syn.GCaMP6f.WPRE.SV40 virus at 50nl/min in dorsal CA1 (−2.1mm AP relative to bregma, +2.0mm ML from bregma, and 1.65mm ventral from the skull surface) using a Nanoject II microinjector (Drummond Scientific). After a week of recovery, mice underwent a microendoscope implantation surgery. A skull screw was fastened to the skull and a 2mm diameter craniotomy was performed above the viral injection site. The cortical tissue above the targeted implant site was carefully aspirated using a 27 gauge blunt needle. Buffered artificial cerebrospinal fluid was constantly applied throughout the aspiration to prevent desiccation of the tissue. The aspiration ceased after partial removal of the corpus callosum and bleeding terminated, at which point a gradient refractive index (GRIN) lens (2mm diameter, 4.79mm length, 0.25 pitch, 0.50 NA, Grintech Gmbh) was stereotaxically lowered to the targeted implant site (−1.35mm DV from the skull surface relative to the most posterior point of the craniotomy). Cyanoacrylate glue and dental cement (Lang Dental) was used to seal and cover the exposed skull, and Kwik-Sil (World Precision Instruments) covered the exposed microendoscope. Carprofen (5 mg/kg) and dexamethasone (0.2 mg/kg) were administered during surgery and for 7 days post-surgery along with amoxicillin (0.25mg/ml) in the drinking water. Three weeks later, animals were again anesthetized and a miniature microscope locked onto an aluminum baseplate was placed on top of the microendoscope. After searching the field of view for in-focus blood vessels and cells, the baseplate was cemented into place and the miniature microscope was unlocked and detached from the baseplate. A plastic cap was locked into the baseplate to prevent debris buildup.

**Wire-free Miniscope:** The wire-free Miniscope is a modified version of the wired Miniscope previously described^21^ with the additional features of being battery powered and logging imaging data onto onboard removable memory. The optics, Delrin housing, and excitation light source are the same between the wired and wire-free Miniscope but the CMOS imaging sensor electronics and DAQ electronics were redesigned and packaged onto a single PCB to enable wire-free operation. All electrical components were chosen to minimize power consumption and weight. The main components of the wire-free PCB are a low power CMOS imaging sensor (E2V JADE EV76C454), ARM Cortex M7 microcontroller (Atmel ATSAME70N21A), and microSD card mount (Molex 0475710001). The PCB also includes necessary voltage regulators, an SD card voltage translation transceiver (TI TXS0206AYFPR), custom excitation LED current driving circuitry, battery connector, and 12MHz oscillator. A single-cell lithium polymer battery (Power Stream GM041215, 45mAH, 1.1g) was attached to the side of the Miniscope and connected to the wire-free PCB during operation. Overall, the wire-free Miniscope weighs between 4-5g depending on the imaging sensor and battery used. Upon power-up, the microcontroller (MCU) accesses configuration information, programmed by custom PC software, in a specific memory block of the microSD card which holds recording parameters such as recording length, framerate, excitation LED intensity, imaging resolution, and imaging exposure length. Once configuration is complete, the MCU waits 5 seconds before beginning recording, at which time the excitation LED and onboard status LED turn on. At the end of the configurable recording duration, the excitation LED and onboard status LED turn off and the MCU terminates data logging onto the microSD card. Offline synchronization of the Miniscope recording with a behavioral camera is achieved through detecting the on and off event of the Miniscope’s status LED.

Pixel values from the CMOS imaging sensor are clocked into a 3 frame circular buffer in the MCU using the MCU’s parallel capture interface and Direct Memory Access (DMA) Controller to minimize processor overhead. Once an imaging frame has been received by the MCU it is written in raw format into incremental memory blocks of the microSD card along with a footer containing timing and error flag information. Wire-free Miniscope data was recorded at 20FPS at a resolution of 320px x 320px using 2x pixel subsampling. Running at this data rate, the entire system consumes approximately 85mW of power during recording.

**Linear Track Imaging:** Prior to the experiment, mice were water restricted and maintained at a body weight of ~85% of initial weight. Mice were handled for five minutes per day for three days and habituated to the miniature microscope for at least ten days with the weight and duration of the sham microscope progressively increasing each day until mice comfortably sustained the full Miniscope for 15 minute sessions. The mice were housed in the same room where the imaging and training sessions were conducted. A wire-free Miniscope, powered by a single-cell, 45mAh lithium polymer battery, was affixed to the mouse’s baseplate. Prior to the beginning of each imaging session, a sample 15 s imaging video was recorded to ensure the proper field of view and the linear track cleaned with 70% ethanol. Data was locally saved onto a microSDHC (SanDisk) card. At the beginning of the linear track recording session, the Miniscope was powered on and the mouse was placed on one end of the 2m track. Mice ran back and forth along the track as experimenters pipetted 10uL water reward at each end of the track. Behavior was recorded with an overhead webcam. After 15 minutes, the microscope powered off and the mouse was taken off of the track and returned to its cage. Imaging sessions were performed every 2 days for 8-10 sessions, followed by shorter interval imaging sessions occurring 24hrs, 6hrs, and 30 mins apart.

**Linear Track Analysis:** Wire-free Miniscope data was extracted from microSD cards and saved as uncompressed 8bit AVI video files for processing and analysis. Animal position was tracked with an overhead behavioral camera that was synchronized offline with the Miniscope data. An FFT based image registration algorithm (courtesy Dario Ringach) was used to correct for frame to frame translational shifts of the brain during animal movement. Constrained non-Negative Matrix Factorization for Endoscopic recordings (CNMF-E)^33^ was used to identify and extract individual neurons’ spatial shapes and fluorescent calcium activity. Fast non-negative deconvolution^34^ was used to deconvolve the extracted fluorescent activity of each neuron. The resulting measure can be interpreted as the probability of a neuron being active at each frame scaled by an unknown constant. For the sake of brevity, we refer to this measure as the “temporal neural activity” of a cell.

Spatial neural activity rates were calculated using 2cm wide spatial bins and a speed threshold of greater than 10cm/s. Temporal neural activity and occupancy of the animal were spatially binned and then smoothed using a Gaussian kernel with sigma = 5cm. The binned neural activity was divided by the binned occupancy to calculate the spatial neural activity rate of each cell.

The information content of a single neuron’s spatial neural activity rate map (in bits)^35^ was defined as:

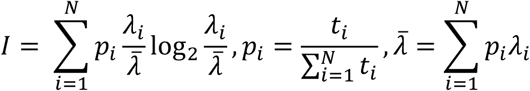

Significance of the spatial information was calculated using a circular trial shuffling procedure. For each trail, the position of the animal was circularly shifted by a random amount and then the spatial activity rate map and information content was calculated across all shuffled trials. This procedure was repeated 500 times to determine a significance measure for each neuron’s spatial activity rate map. The significance measure is the percentile of the true information content value within the distribution of the information content values from the 500 randomly shuffled sessions.

Stability of a single neuron’s spatial activity rate map was calculated by taking the Fisher Z-score of the Pearson correlation coefficient between the spatial activity rate maps at two time points. Within session stability was calculated by averaging the stability measure of odd vs even trials with the stability measure of first half vs second half of trails. The same circular trial shuffling procedure mentioned above was applied to calculate significance of within session stability.

The Population Vector Overlap (PVO) for a population of N neurons over a track length L was defined as^35^:

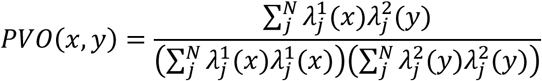

Where 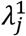 and 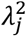 are the spatial activity rate maps of neuron *j* at two different time points. PVO curves were generated by calculating the mean PVO value for all PVO bins with a Δx difference in positions between the two time points being compared. Δx was varied from 0 to 150cm to generate the full PVO curve.

**Place Cell Criteria:** For a cell to be classified as a place cell it must satisfy the three following criteria: 1) The neuron’s information content is above chance (P<0.05) based on the circularly shuffled distribution, 2) the neuron’s within session stability is above chance (P<0.05) based on the circularly shuffled distribution, 3) the neuron’s spatial activity rate map has consecutive bins spanning at least 10cm with activity rates in at least the 95^th^ percentile of binned activity rates of circular trial shuffled spatial activity rate maps.

**Matching Cells Across Sessions:** First, field-of-view (FOV) shifts between sessions were corrected by performing rigid image registration between the mean frame of two sessions. For all pairs of cells across two sessions, the centroid distance and Pearson correlation of spatial footprints was calculated. Those pairs whose centroid distance was less than or equal to 4 pixels (~7um) and whose spatial footprint correlation was greater than or equal to 0.6 we counted as the same cell. More information can be found in Extended Data 4.

**Bayesian Decoder:** A Bayesian decoder was constructed to evaluate how well neural activity estimated the animal’s location based on an independent training set of neural activity (Extended Data 10). For within session decoding, the training data set was generated from 3/4^th^ of the trials evenly distributed across the session and then the decoder was applied to the remaining 1/4^th^ of interspersed trials. For across session decoding, the training data set was generated from the entire session of the first time point and then the decoder was applied to the entire session of the second time point. All decoders used the top 100 neurons with the highest within session stability in the training session and that showed up in both training and decoding sessions.

Temporal neural activity (deconvolved fluorescence activity) was first binarized for both the training data set and decoding data set. For each neuron, time points with zero probability of neural activity were kept at zero, all other time points were set to one. Data points where the animal was in a reward zone or when the animal’s speed was under 10cm/s were excluded from the training and decoding data sets. For the training data set, the binarized spatial neural activity rate per trial per neuron was calculated. Binarized neural activity and occupancy were spatially binned along the length of the track, smoothed using a Gaussian kernel (sigma = 5cm), and then the binned activity was divided by the binned occupancy. For each neuron in the training set, the mean and standard deviation (SD) of the per trial binarized spatial neural activity rate were calculated per spatial bin. Bins with SD below that neuron’s mean per bin SD were raised to the mean SD. The training data set for each neuron resulted in a Guassian distribution for each spatial bin along the track which was used as the probability function in the decoder.

To estimate the animal’s position 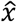 at time *t* using the neural activity of *N* neurons present in both the training data set and decoding data set the decoder was defined as

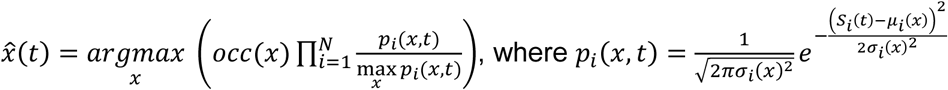

*occ(x)* is the occupancy per spatial bin *x* from the training data set. *μ*_*i*_(*x*) is the mean and *σ*_*t*_*(x)* is the SD of the binarized neural activity rate of at spatial bin *x* of neuron *i* from the training data set. *S*_*i*_*(t)* is the mean binarized neural activity rate of neuron *i* from the decoding data set over time time *t* ± Δt, where Δt *=* 0.125*s*.

**In vivo Electrophysiology Surgeries:** For in vivo electrophysiology recordings, all mice underwent two stereotaxic surgeries. First, a stainless steel headbar was fixed onto the mouse’s skull. The mouse’s skull and headbar were each stereotaxically aligned and the headbar was lowered near the skull and fixed onto the mouse’s skull using cyanoacrylate glue and dental cement. Dental cement was build up around the perimeter creating a ring around the exposed skull, and filled with Kwik-Sil. A final layer of dental cement was applied above the Kwik-Sil and dental cement perimeter. Carprofen (5 mg/kg) was administered both during surgery and for 2 days post-surgery along with amoxicillin (0.25mg/ml) for 7 days post-surgery.

One day prior to recording, a craniotomy and ground implantation was performed. First, the most superficial layer of dental cement was drilled off and the Kwik-Sil was removed. Mice were stereotaxically aligned and an approximately 2mm × 1mm craniotomy was performed over the target recording site (−2.1mm AP, +/− 1.5mm ML). A second craniotomy was performed above the cerebellum, and a silver reference wire (Warner Instruments) was slipped between the skull and dura and fixed with dental cement. The exposed skull was covered with Kwik-Sil and the mouse was allowed to recover overnight.

**VR Training and Recording:** One week after headbar surgery, mice were water restricted and maintained at a body weight of ~85% of initial weight. Mice were handled for five minutes each day for three days and habituated to head-fixation on a spherical treadmill, locked to rotate only on 1 axis, for 3-5 days. Next, mice were trained to lick from a water tube by allowing a 5 μL water reward during each lick. Once mice reliably licked from the water tube, they were immediately trained to run along the virtual reality linear track. The virtual reality apparatus was similar to previously described^36^ and consisted of a projector (Epson), a flat mirror, a circular mirror, and a toroidal screen that surrounded the animal (Fig 3A), and the display was warped to match the curved screen with NTHUSIM software. The virtual linear track was created with Platinum Arts Studio Sandbox and modified to trigger water delivery and record location and reward data through a custom data acquisition device. Mice were trained to run forward to reach the end of the track and receive a water reward. Mice were first trained on a short track until they completed >100 trials within 45 minutes, and were then moved to a longer track. This was repeated until animals consistently ran on the longest track (total training was approximately 5-20 days). A 5uL water reward was dispensed at the end of the virtual track through a metal tube controlled by a solenoid valve (Lee Company). After five consecutive and consistent training sessions, mice were deemed ready for recording. On the day of the recording, mice were head-fixed onto the treadmill, the Kwik-Sil covering the craniotomy was removed and replaced with buffered ACSF and the mouse was aligned to the micromanipulator. A 128 channel silicon probe^37^ was slowly lowered into the hippocampus and the surface of the exposed brain was covered with mineral oil. After at least 60 minutes to allow the brain to settle, the recording began and continued for 180 minutes.

Silicon probe data was collected similar to previously described^37^. Signals were amplified (gain=200), filtered (0.1-8000hz), and multiplexed (32 channels per output wire) by custom headstages using integrated electronic circuits (Intan Technologies, RHA-2164B). Analog signals were simultaneously digitized on National Instruments data acquisition devices (USB-6356) using Labview software (National Instruments). Each channel was sampled at a rate of 25kHz. Licking and reward times were also recorded simultaneously on the same data acquisition device. Licking behavior was monitored with an IR beam lickometer (Island Motion). Trial start times, position, and reward times were also saved through a custom recording software connected via Ethernet to the virtual reality software.

**Electrophysiology Analysis**: All data analysis was carried out with custom Matlab scripts. Data files were first demultiplexed into continuous signals. All analysis was restricted to times of locomotion and interictal spikes were removed. For LFP analysis, signals were bandpass filtered using zero-phase digital filtering and phase relationships were measured using the Chronux toolbox^38^. Sublayers of the hippocampus were identified from stereotaxic coordinates and electrophysiological markers including peak theta and ripple power^39^, theta phase shifts^39^, gamma coherence^40^, dentate spike phase reversal^41,42^ and the density of single units. A subset was confirmed with post hoc histology. Spike sorting was performed using PyClust software^43^. All channels from each sublayer of the hippocampus were background subtracted, bandbass filtered 600-6000hz, and putative spikes were extracted from signals at least 4 standard deviations from mean. Using peak amplitude, trough amplitude, and principal components, single units were isolated by manual clustering. Putative interneurons were identified based on firing rate, mean autocorrelogram, and complex spike index^42–44^. For phase locking analysis, LFPs were filtered and the Hilbert transform was used to determine the phase of each spike.

For each interneuron, phase angles from each spike were then used to calculate a Raleigh’s r-value and mean phase and significance of phase locking was tested with a Raleigh’s test for nonuniformity. Group differences in the distribution of preferred phases was assessed with the Kuiper test using the CircStats toolbox in Matlab^45^.

**Basic Structure of the CA1 network model:** Simulations of place cell formation were performed using a small CA1 network consisting of biophysical neurons (Hodgkin-Huxley formalism) with reduced morphology^46^. The network model was based on structure and connectivity previously described^47–50^ and consists of 80 principal neurons (pyramidal cells - PCs) and six types of interneurons. These include two axoaxonic cells (AACs), eight basket cells (BCs), two bistratified cells (BIS), two oriens-lacunosum moleculare cells (OLMs), one VIP+/CCK+ cell and four VIP+/CR+ cells. All single cell models were validated against experimental evidence with respect to passive and active properties. External input to the network was provided by entorhinal cortex layer III (EC LIII), CA3 Schaffer Collaterals and Medial Septum (MS). EC LIII and CA3 provided the network with excitation, while afferents from MS contacted only interneurons with inhibitory synapses to generate the theta cycle. All excitatory inputs were grid-like inputs and EC LIII inputs preceded CA3 inputs by 95 ms. Principal neurons also received excitatory background activity on their dendrites, in the form of AMPA, NMDA, GABAA and/or GABAB synaptic currents. All interneurons received excitation via AMPA synapses, whereas PCs received excitation via AMPA and NMDA receptors in their apical and basal dendrites. GABAA was the main inhibitory synapse; however inhibitory synapses in pyramidal dendrites had both GABAA and GABAB receptors. The hallmark properties of the network were: (a) the formation of place cells along a 1-D linear track^51^, (b) the increased number of place cells encoding reward location (enrichment) in the presence of a simulated reward zone^52^ and (c) the decrease in enrichment upon removal of VIP+^46^.

**Place cell formation and simulated experiments:** Grid-like inputs were simulated as simple summation of three sinusoidal waves^53^. Specifically, a single place field emerged from the convergence of eight grid cells differing in their size and phase. All PCs in the network received a convergent octal of inputs. However, a portion of PCs had reduced synaptic weights in order to reproduce silent place cells reported in CA1^54^. To cover the linear track (200 cm in length) with place cells, we simulated 21 theoretical place cells located from 0 to 200 cm with a step of 10 cm. Thus, 168 inputs from EC LIII and 168 from CA3 provided the network with external excitatory input. Each PC received a convergent octal of grid-like inputs to its distal apical, medial apical and basal dendrites. If a PC was a silent place cell, its synaptic weights from EC LIII and CA3 were set to 10% of their control values (i.e., those of active place cells).

To simulate changes observed in epileptic animals^9,10,55^, the total number of SOM+ (OLM) interneurons was reduced by 50%, that of PV+ (AAC, BC, BSC) interneurons was reduced by 30%, and the EC LIII and CA3 inputs were desynchronized by ~ ±20 ms. For desynchronization, spike timings in CA3 were shifted by a normal distribution with mean *μ =* 0 *ms* and standard deviation *σ = 20 ms.* Simulations with combinations of the aforementioned manipulations were also performed.

**Analysis of modeling data:** For each pyramidal cell, we calculated its information content, stability, field size and peak firing rate. Information content and stability were calculated similar to experimental place cell data (see Linear Track Analysis above). The field size was defined as the number of consecutive bins in both directions from the peak, in which the firing rate exceeded 10% of the peak (maximum) firing rate. Occupancy information was circularly shuffled by a random amount for each of the trials, and the information content and stability index of the resulting rate map was computed. This procedure was repeatedly and independently applied to generate two null distributions of 500 information content values and 500 stability index values. A cell was considered to have significant spatial information content and/or stability index if the information content and/or stability index of its true rate map exceeded that of 95% of the values in the corresponding null distribution. A place cell was defined as a cell which had field size above 8cm and below 50cm and its peak firing rate was greater than 2 Hz.

An “animal” represented a network with a slightly different connectivity profile (connection probabilities were kept the same). For each of our “animals” we had several repetition trials (50 trials) where the input was slightly different due to noise but the network connectivity remained the same. All simulations were performed in a high performance computing cluster with 312 cores under the CentOS 6.4 Linux operation system. The CA1 network and all individual neuronal types were simulated in the NEURON simulation environment^56^, whereas the grid-like inputs, the background noise and the analysis were constructed using python (2.7.13) custom-made software.

### Statistics

Statistics were performed using Matlab (Mathworks), SPSS (IBM), and GraphPad Prism software. Statistical significance was assess by two-tailed unpaired Student’s t-tests, Mann-Whitney U test, one-way ANOVA, or two-way ANOVA where appropriate. Normality was assessed using Kolmogorov-Smirnov test for normality. For non-normal distributions, nonparametric Mann-Whitney tests were performed. Significant main effects or interactions were followed up with post hoc testing using Bonferroni corrections where applicable. Significance was declared at P<0.05, with a precise P value stated in each case. For circular tests, group differences in the distribution of preferred phases was assessed with the Kuiper test. Due to poor behavior (<10 trials run), one session from one Control mouse and four sessions from one Epileptic mice were removed from Figure. 1 and 2.

### Data Availability

The experimental data that support the findings of this study are available from Peyman Golshani. The software/codes related to the CA1 network model and its analysis are available from the Poirazi lab. The model will also be available at www.dendrites.gr and ModelDB, following publication.

### Wire-Free Miniscope Availability

All design files, software, parts lists, and tutorials necessary to build and implement the wire-free Miniscope will be available at www.miniscope.org.

## Acknowledgements

We thank Katie Maguire, Jerry Lou, Aria Fariborzi, Naina Rao, Justin Daneshrad, Shayan Ghiaee, Ryan Manavi, Cynthia Araradian, Celina Yang, Melissa Song, My La-Vu, Brandon Wei, Chengbin Zhou, Ashley Meyer, Hesper Chen, James Davis, Nora Abduljawad, John Hodson, Iris Bachmutsky, Leeor Zilbermintz, Haleh Karbasforoushan, Jonathan Friedman, and Theunis Kotze for excellent technical assistance and help with experiments. We thank Naina Rao for assistance with graphical design. This work was supported by VA Merit Award 1 I01 BX001524-01A1, U01 NS094286, R01MH101198, R01 MH105427, U54 HD87101, R01NS099137, NSF Neurotech Hub 1700408 to P.G.; David Geffen School of Medicine Dean’s Fund for development of open-source miniaturized microscopes to B.S.K., A.J.S. and P.G.; Cellular Neurobiology Training Grant T32 NS710133 and Epilepsy Foundation Postdoctoral ResearchTraining Fellowship to T.S.; Neurobehavioral Genetics Training Grant T32 NS048004 and Neural Microcircuits Training Grant T32 NS058280 to D.A.; National Research Service Award F32 MH97413 and Behavioral Neuroscience Training Grant T32 MH15795 to D.J.C.; DP1 MH104069 to B.S.K.; McKnight Technological Innovations in Neuroscience Award to S.C.M.; Dr. Miriam and Sheldon G. Adelson Medical Research Foundation to A.J.S.

## Author Contributions

TS, PG designed the calcium imaging and electrophysiology experiments. TS, DA, DJC, BSK, AJS, PG developed the wire-free Miniscope. DA designed and built the wire-free Miniscope. TS, DA, DJC, CL tested the wire-free Miniscope. TS, CL, DC performed calcium imaging experiments. TS, DA analyzed calcium imaging experiments. TS, MJ, CCK, MS, PG designed the virtual reality apparatus. TS, SEF, KC, MJ, CCK performed electrophysiology training and recording. TS, KIB, SCM, PG built the electrophysiology system. TS, JT analyzed electrophysiology data. TS, KC, CCK performed spike sorting of single units. TS, DA, DJC, SC, PP, PG designed modeling experiments. TS, SC, PP performed and analyzed modeling experiments. TS, DA, DJC, SC, PP, PG wrote the manuscript and all authors edited the manuscript.

## Author Information

Correspondence and requests for materials should be addressed to.

Correspondence and requests for materials related to modelling experiments only should be addressed to.

**Extended Data 1.**
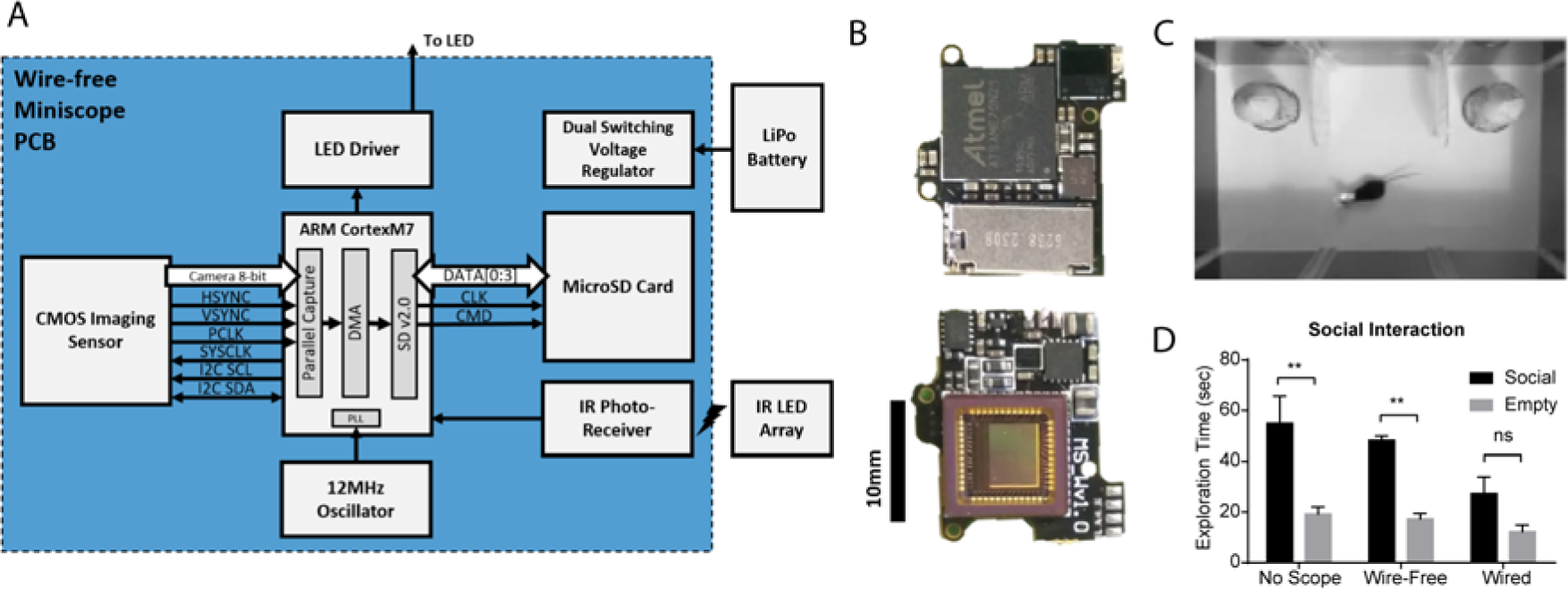
Wire-free Miniscope design and behavior. A. Schematic of wire-free Miniscope printed circuit board (PCB). B. Picture of top and bottom of wire-free sensor board. This camera sensor board attaches to the Miniscope body, which contains all optic components. C. The wire-free Miniscope can be used to probe naturalistic behaviors including social interaction. We tested social behavior with the wire-free Miniscope compared to no Miniscope or a wired version. D. During a social interaction test mice with either no Miniscope or the wire-free Miniscope chose a social cup over an empty cup, while mice with a wired Miniscope showed no significant preference (n=3 per group, Unpaired t-tests, No Scope: P=0.03, Wire-Free: P<0.001, Wired: P=0.10). Error bars represent 1 S.E.M. *P<0.05, **P<0.01.

**Extended Data 2.**
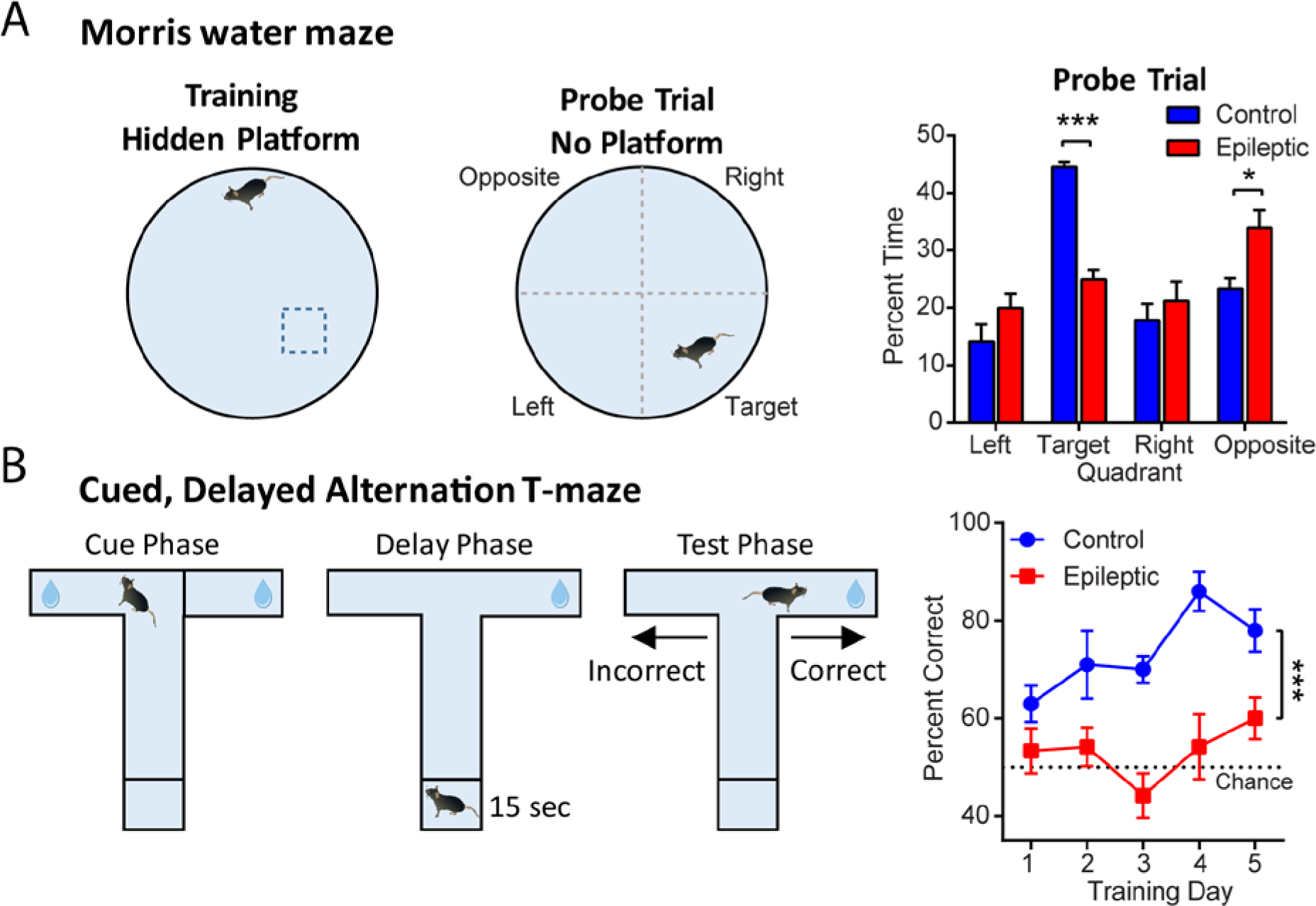
Spatial memory deficits in epileptic mice. A. Morris water maze task. Mice were trained to find a hidden platform for 6 days at random start locations. The next day, mice were given a probe trial to assess learning. On the probe trial, control mice spent more time in the target quadrant than epileptic mice, while epileptic mice spent more time in the opposite quadrant (n=6 Control, n=5 Epileptic, F_GroupXQuadrant_(3,36)=14.6, P<0.001, Training Quadrant Post-hoc: P<0.001, Opposite Quadrant Post-hoc: P<0.05). B. Mice were trained on a cued, delayed alternation T-maze task for 5 days. On each trial, animals were cued to one direction (Cue Phase), returned to the start position for a 15 second delay (Delay Phase), and then released for the Test Phase. If the animal went to the opposite side it received a water reward. Epileptic mice performed worse than control mice (n=5 Control, n=6 Epileptic, FGroup(1,45)=44.98, P<0.001). Error bars represent 1 S.E.M. *P<0.05, ***P<0.001.

**Extended Data 3.**
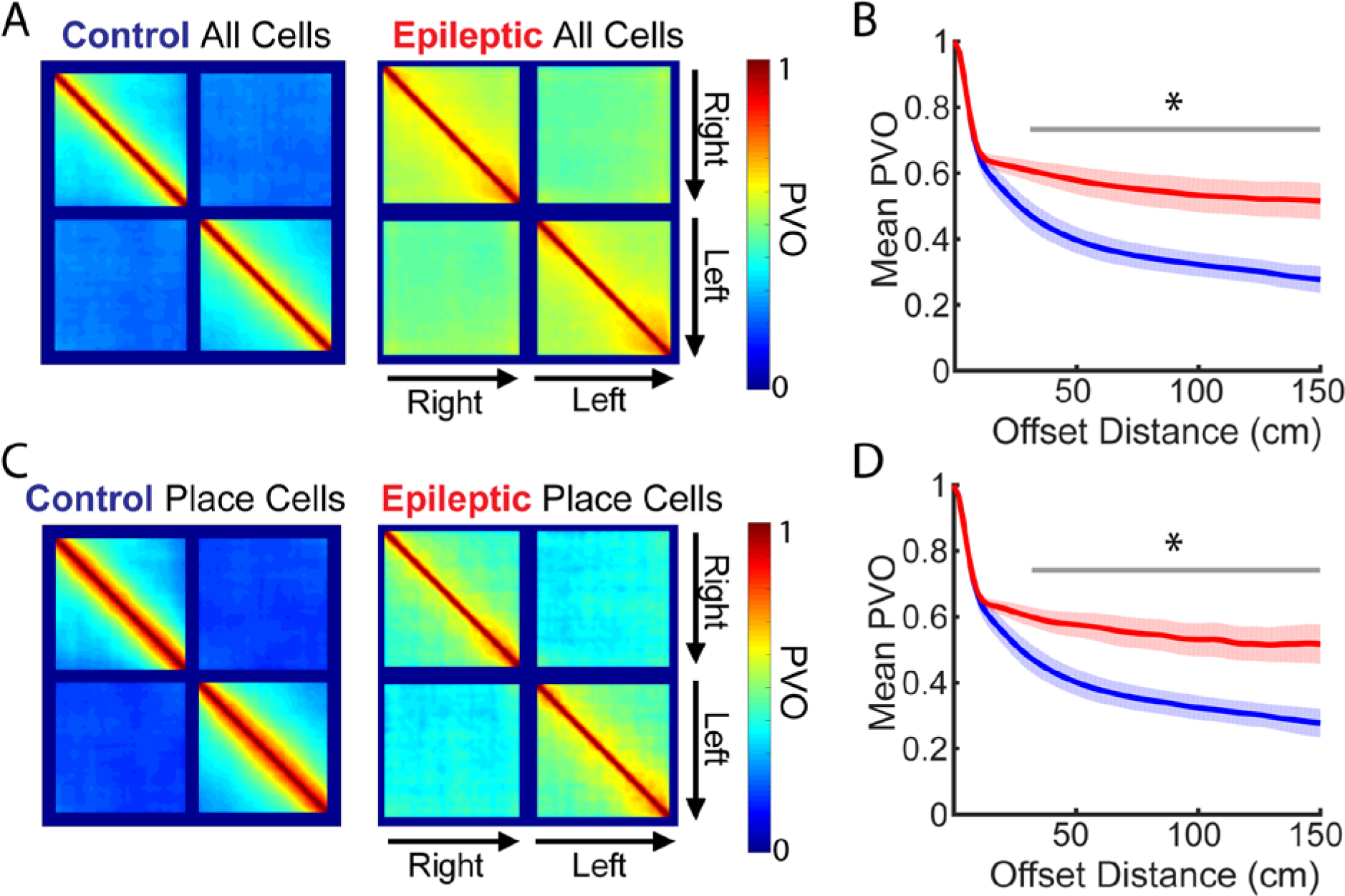
Within Session PVO. A. Population Vector Overlap (PVO) of all cells in Control (left) and Epileptic (right) mice across all sessions recorded. B. Mean PVO as a function of offset distance (from the diagonal) across animals. Epileptic mice had higher PVO (i.e., less distinct firing patterns) across distances of 30-150cm (FGroup(1,8)=10.39, P=0.01, post hoc P<0.05 for each bin from 30-150cm). C. Population Vector Overlap (PVO) of place cells in Control (left) and Epileptic (right) mice across all sessions recorded. D. Mean PVO as a function of offset distance (from the diagonal) across animals. Epileptic mice had higher PVO (i.e., less distinct firing patterns) across distances of 32-150cm (FGroup(1,8)=10.19, P=0.01, post hoc P<0.05 for each bin from 32-150cm). Error bars represent 1 S.E.M. *P<0.05.

**Extended Data 4.**
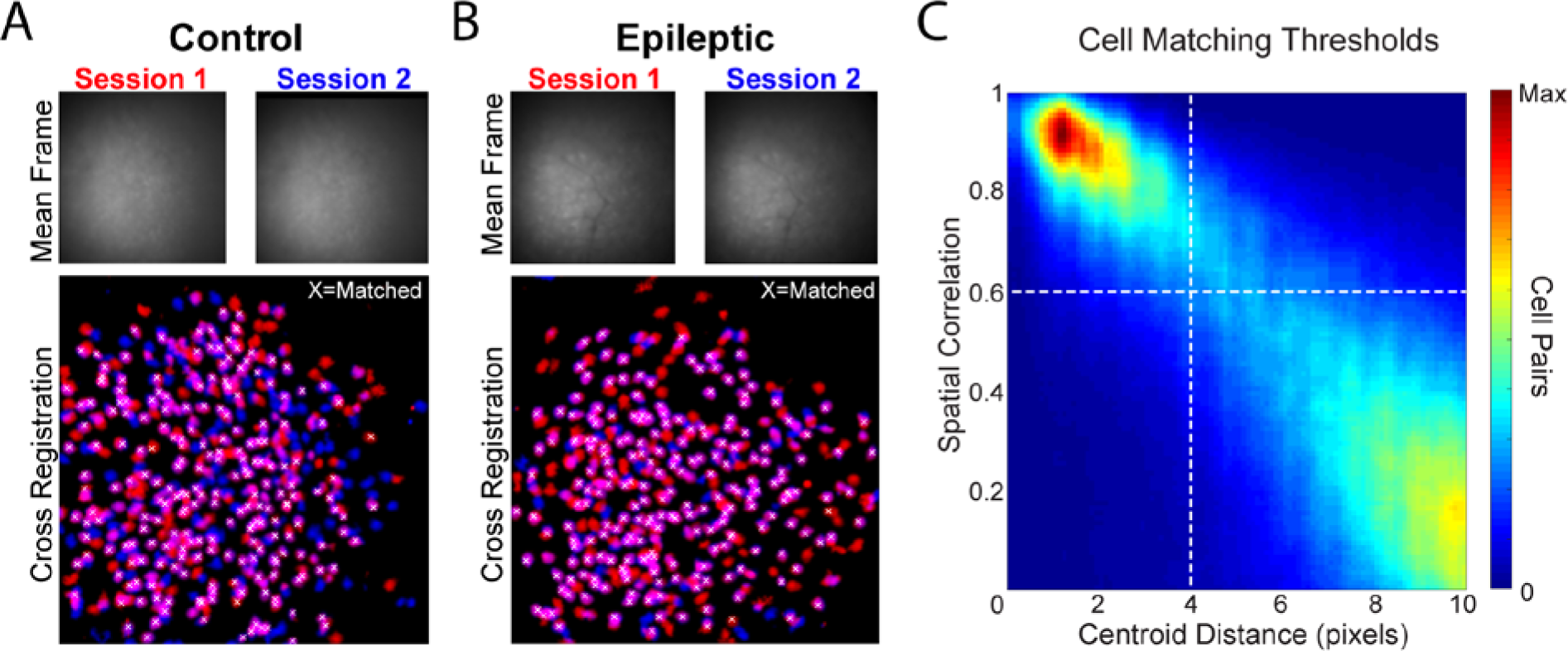
Cross Registration. A-B. Example cross registration in a Control (A) and Epileptic (B) mouse. Top, Aligned mean frame from each session (~550um × 550um). Bottom, overlaid cells from each session with white X indicating matched cells. C. Spatial correlation and centroid distance were calculated for all cell pairs. Dotted lines indicate thresholds used as matching criteria. All matched cells had spatial correlation ≥ 0.6 and centroid distance ≤ 4 pixels (~7um).

**Extended Data 5.**
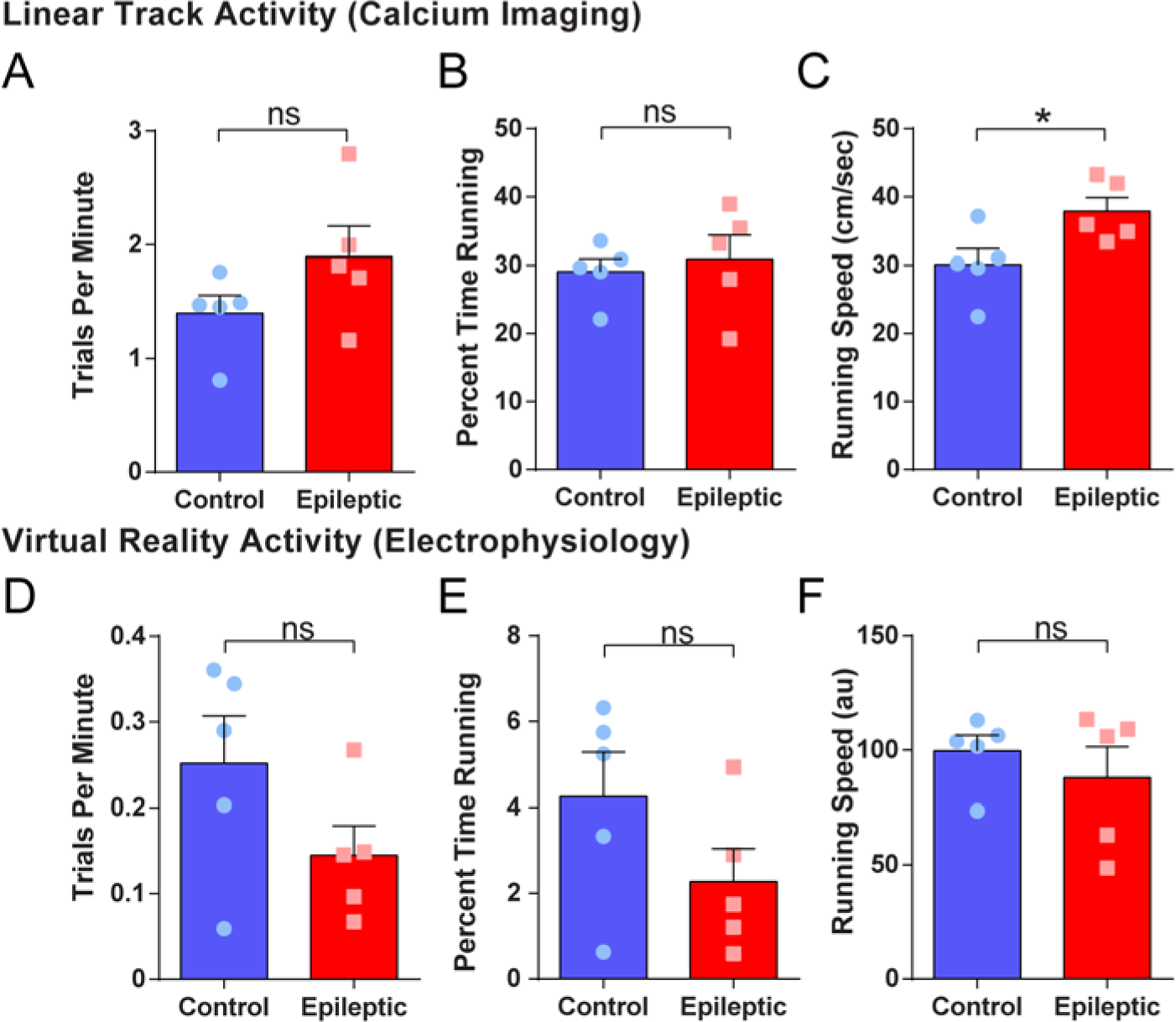
Running activity during calcium imaging and electrophysiology. A-C. Activity during calcium imaging on the linear track. No differences were found in the number of trials per minute (n=5 per group, Mann-Whitney test, P=0.15) or percent time running (n=5 per group, Unpaired t-test, P=0.64). Epileptic mice ran faster than control mice on the linear track (n=5 per group, Unpaired t-test, P=0.03). D-F. Activity during virtual reality recordings. No differences were found in trials per minute (n=5 per group, Unpaired t-test, P=0.16), percent time running (n=5 per group, Unpaired t-test, P=0.14), or running speed (n=5 per group, Mann-Whitney test, P=0.94). Error bars represent 1 S.E.M. *P<0.05.

**Extended Data 6.**
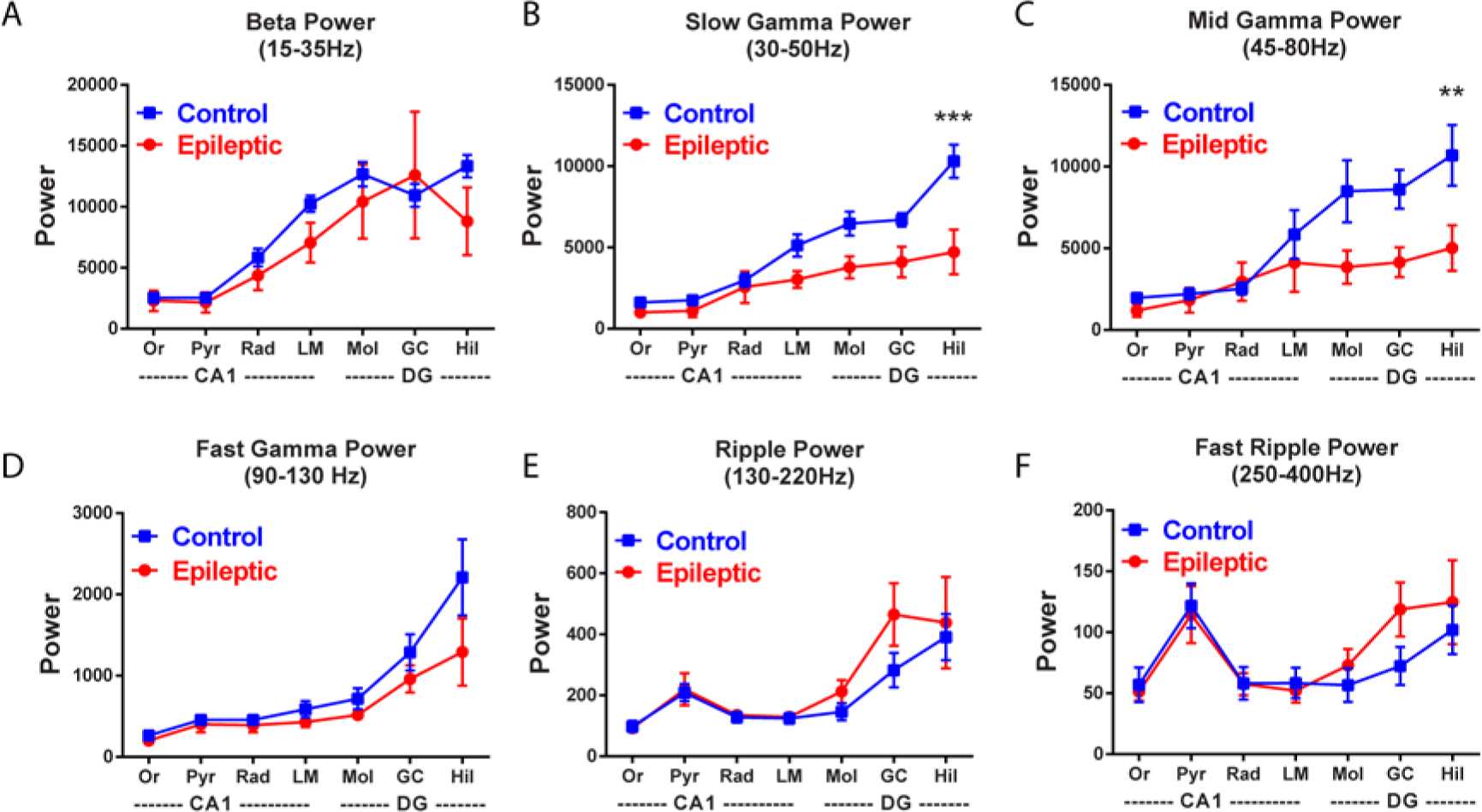
Local field potential power across frequencies. A-F. LFP power in beta, slow gamma, mid-gamma, fast gamma, ripple and fast ripple frequencies throughout CA1 and DG. No differences were found in beta (F_GroupXRegion_(6,48)=1.23, P=0.31, FGroup(1,8)=0.47, P=0.51), fast gamma (F_GroupXRegion_(6,48)=1.88, P=0.10, FGroup(1,8)=2.03, P=0.19), ripple (F_GroupXRegion_(6,48)=0.93, P=0.48, FGroup(1,8)=0.67, P=0.44), or fast ripple (F_GroupXRegion_(6,48)=1.38, P=0.24, FGroup(1,8)=0.24, P=0.64) power. Epileptic mice had reduced slow gamma and mid-gamma power in the hilus of DG (Slow Gamma: F_GroupXRegion_(6,48)=6.76, P<0.001, F_Group_(1,8)=7.31, P=0.03, Hilus: P<0.001; Mid-gamma: F_GroupXRegion_(6,48)=4.49, P=0.001, F_Group_(1,8)=3.48, P=0.09, Hilus: P<0.01). n=5 per group for all graphs. Error bars represent 1 S.E.M. **P<0.01, ***P<0.001

**Extended Data 7.**
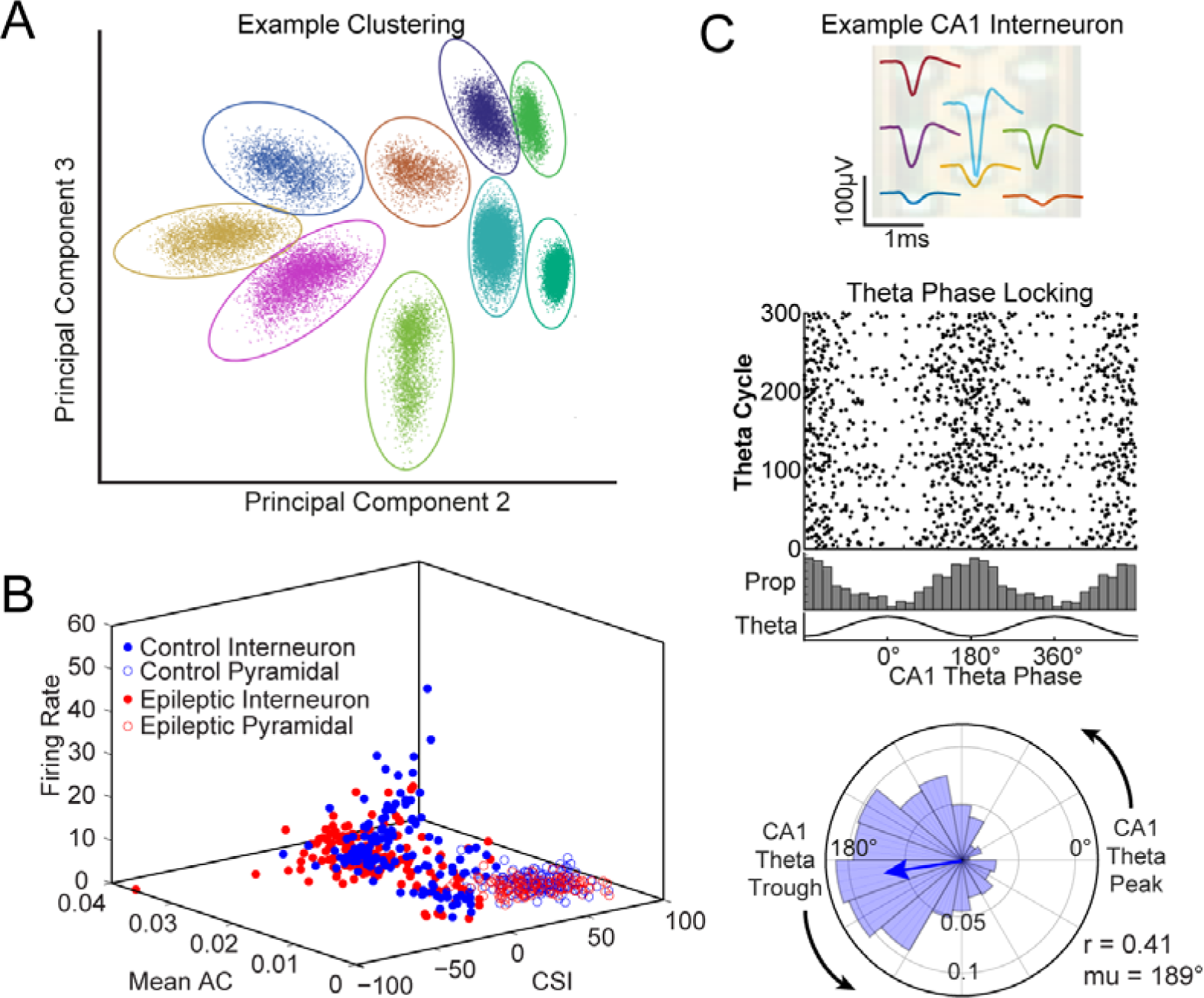
Unit clustering and phase locking. A. Example clustering of single units using principal components, peak amplitude, and trough amplitude. B. Units were characterized as putative interneurons based on complex spike index (CSI), mean autocorrelogram (Mean AC), and mean firing rate. C. Example phase locking in a CA1 interneuron. Top, mean waveforms from each recorded channel within the pyramidal layer. Middle, spike raster of 300 theta cycles during running, with the proportion of spikes occurring during each phase of theta. Bottom, Rose plot of firing phase relative to theta. This cell has a modulation index (r-value) of 0.41 and preferred firing phase (mu) of 189°.

**Extended Data 8.**
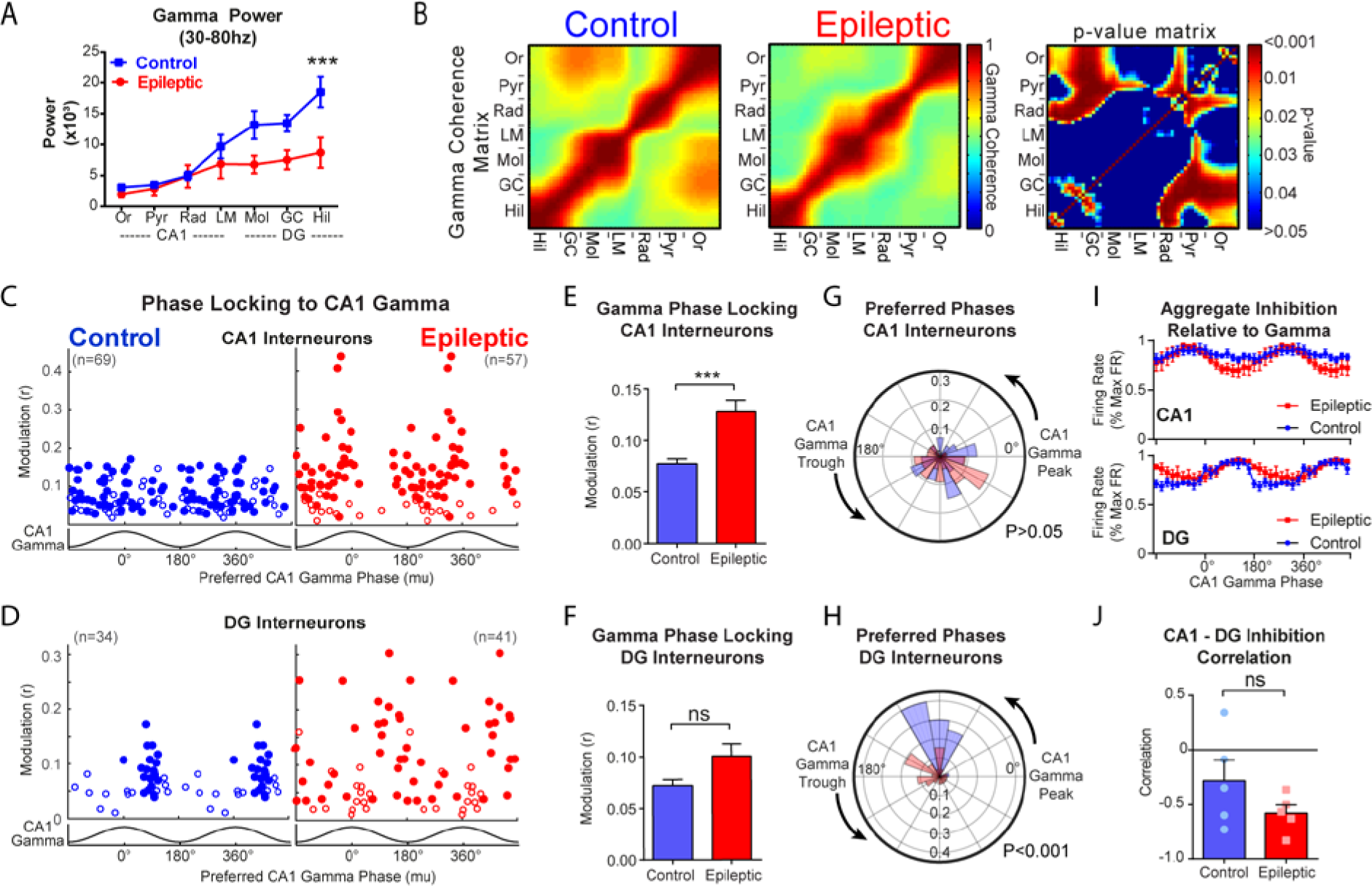
Altered gamma synchronization in epileptic mice. A. Gamma power throughout CA1 and DG. Epileptic mice had reduced gamma power in the hilus of dG (F_GroupXRegion_(6,48)=5.26, P<0.001, Hilus: P<0.001). B. Gamma coherence between each channel pair along the recording probe in control (left) and epileptic (middle) mice. Right, p-value matrix for each pair of recording sites. C. Phase locking to CA1 gamma for each interneuron in CA1 of control and epileptic mice. Each dot represents one interneuron and the data is double plotted for visualization. Closed circles were significantly phase locked (Raleigh test) and open circles were not. D. Phase locking to CA1 gamma for each interneuron in DG of control and epileptic mice. E. Mean phase modulation (r) of CA1 interneurons to CA1 gamma. Interneurons in epileptic mice had increased phase modulation (n=69 Control cells, n=57 Epileptic cells, Mann-Whitney test, P<0.001). F. Mean phase modulation (r) of DG interneurons to CA1 gamma. Interneurons in epileptic mice had increased phase modulation (n=34 Control cells, n=41 Epileptic cells, Mann-Whitney test, P=0.42). G. Rose plot of preferred firing phases of significantly phase locked CA1 interneurons. There were no differences between interneurons in control and epileptic mice (Kuiper circular test, P>0.05). H. Rose plot of preferred firing phases of significantly phase locked DG interneurons. The distribution of preferred phases in control and epileptic mice were different (Kuiper circular test, P<0.001). I. Aggregate inhibition of all interneurons in CA1 and DG relative to CA1 gamma phase (CA1: F_GroupXPhase_(17,136)=1.22, P=0.25, DG: F_GroupXPhase_(17,136)=2.196, P<0.01). M. Correlation between CA1 and DG aggregate inhibition relative to gamma. No difference was found between epileptic and control animals (Unpaired t-test, P=0.19). All data in this figure came from 5 animals per group. Error bars represent 1 S.E.M. *P<0.05, ***P<0.001, ns: not significant

**Extended Data 9.**
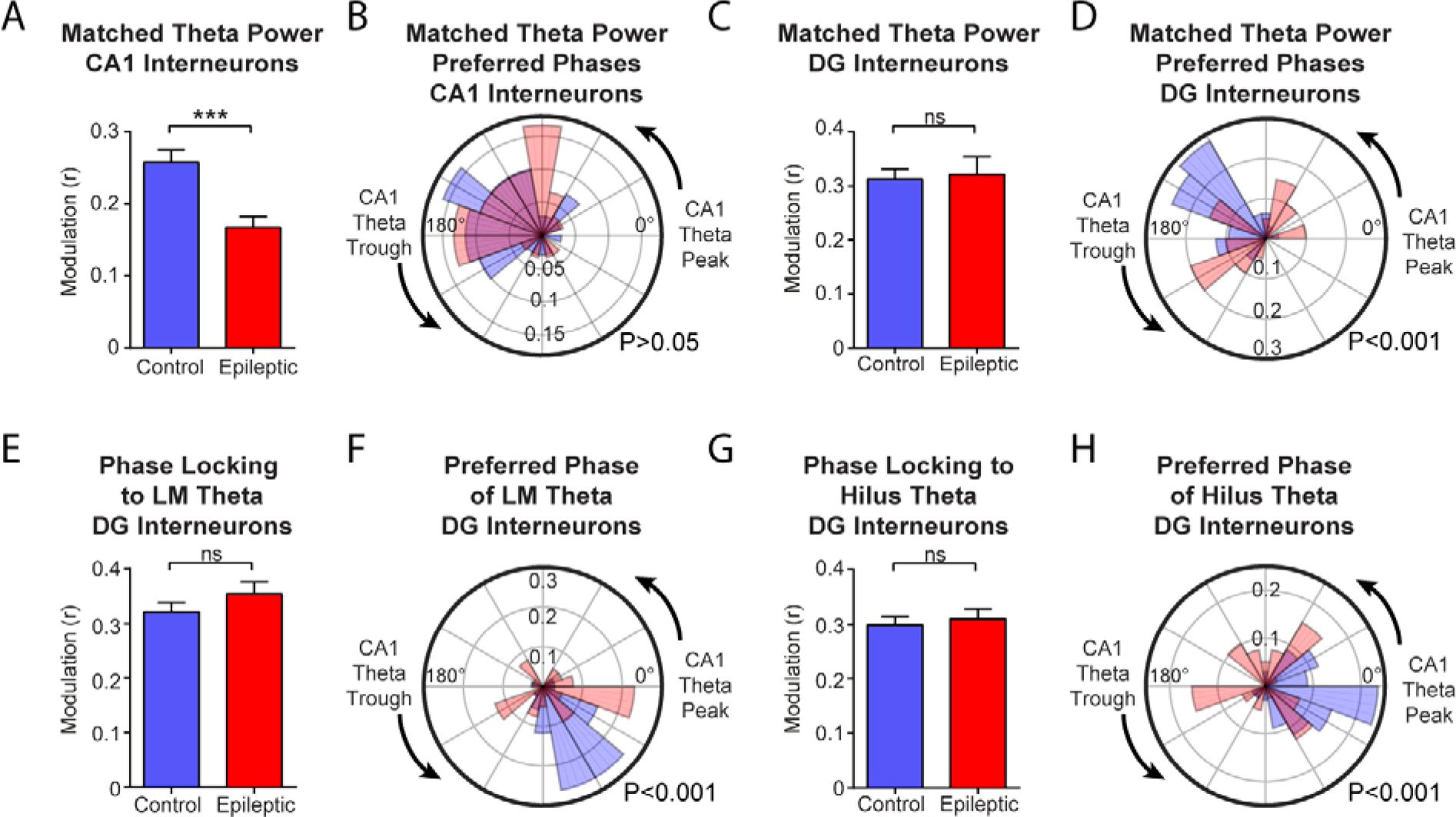
Theta phase locking changes are not caused by decreased power or specific to reference location. A-D. Theta cycles were subsampled to match power between control and epileptic mice. This did not change the results as CA1 interneurons in epileptic mice were less phase locked than in control mice (A; n=71 Control cells, n=34 Epileptic cells, Unpaired t-test, P<0.001), while their preferred phase was not different (B; Kuiper circular test, P>0.05). There were no differences in the modulation of DG interneurons (C; n=34 Control cells, n=22 Epileptic cells, Unpaired t-test, P=0.80) but there were differences in the preferred phase of DG interneurons (D; Kuiper circular test, P<0.001). E-F. Phase locking of dG interneurons to theta in the lacunosum moleculare. No difference was found in modulation (E; n=34 Control cells, n=41 Epileptic cells, Unpaired t-test, P=0.25) but the distributions of preferred phases differed between the groups (F; Kuiper circular test, P<0.001). G-H. Phase locking of DG interneurons to the theta in hilus of DG. No difference was found in modulation (G; n=34 Control cells, n=41 Epileptic cells, Unpaired t-test, P=0.64) but the distributions of preferred phases differed between the groups (H; Kuiper circular test, P<0.001). Error bars represent 1 S.E.M. ***P<0.001.

**Extended Data 10.**
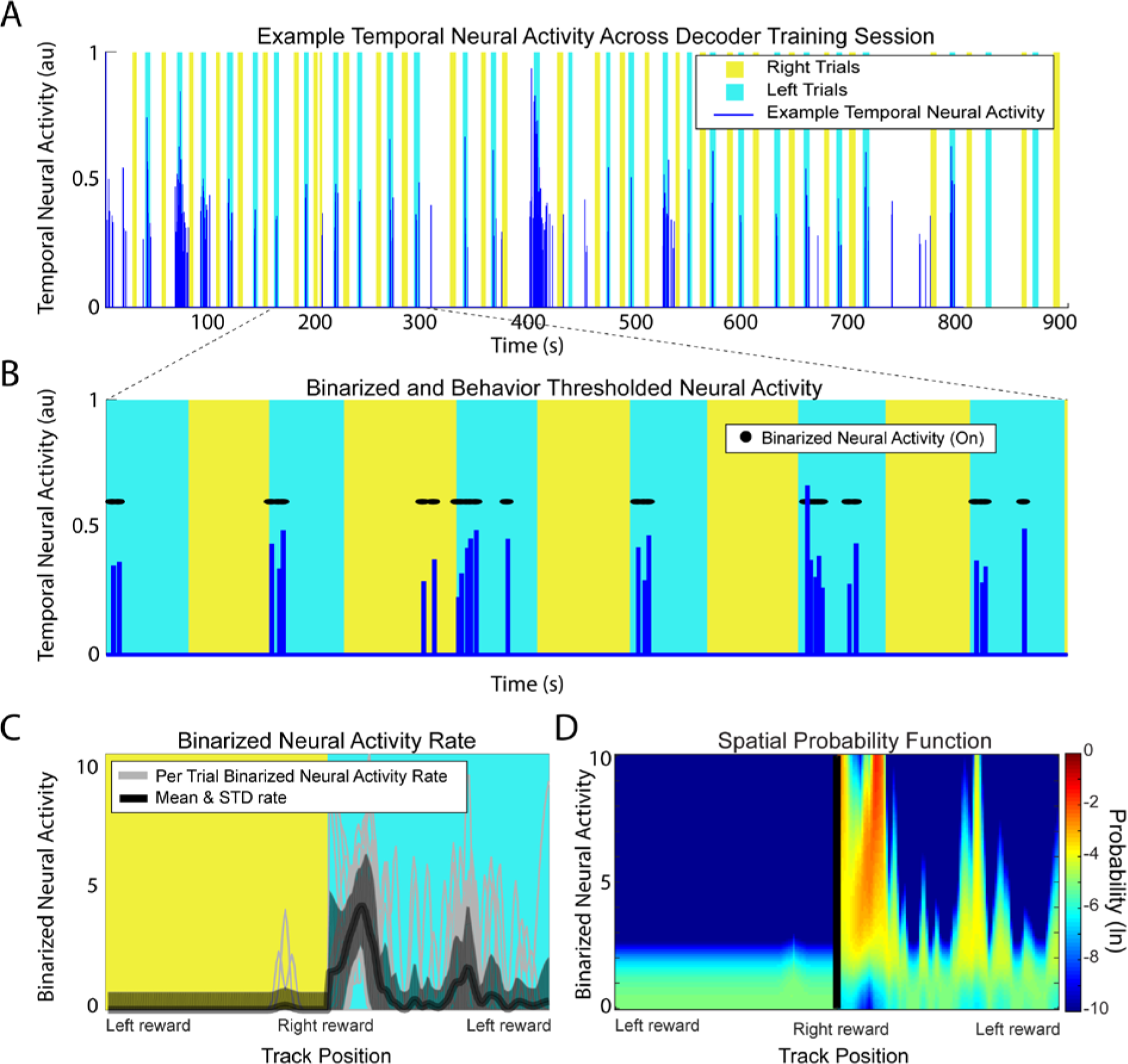
Example processing of one training session neuron for Bayesian decoding. A. For each frame, temporal neural activity is calculated and classified into rightward trials, leftward trials, or non-trial times. B. Temporal neural activity is binarized and non-trial times are removed. C. The per trial binarized neural activity rate is calculated for rightward and leftward trials. D. The spatial probability function is constructed for each cell. A Gaussian distribution is first generated for each spatial bin using the mean and standard deviation of the binarized neural activity rate. The overall distribution is normalized across binarized neural activity rate rows and this data is entered into the Bayesian decoder for each cell. See Supplementary Video 2 for example decoding across all cells.

